# Remote ischemic conditioning modulates inflammatory response and metabolic pathways

**DOI:** 10.1101/2023.08.04.551927

**Authors:** Coral Torres-Querol, Reinald Pamplona, Gloria Arqué, Francisco Purroy

## Abstract

Remote ischemic conditioning (RIC) is an endogenous procedure that reduces ischemic injury by repeated transient mechanical obstruction of vessels at a remote limb. However, the specific mechanism of this protective phenomenon remains incompletely understood. We aimed to study perturbations in the brain and plasma metabolome following RIC as well as to identify potential novel inflammatory cytoprotective targets.

A mouse model of transient focal cerebral ischemia by compressing the distal middle cerebral artery was used. Multiplex cytokine assay was performed in plasma samples. Blood plasma and brain samples were collected and metabolomes analyzed using non-targeted LC-MS.

The analysis revealed a moderate impact on the brain metabolome compared to circulatory metabolites following RIC intervention. Interestingly, 3 plasma metabolites, Cer(42:3), HexCer(36:1) and TG(28:0), stood out as highly significantly upregulated. Moreover, RIC applied during the ischemia (RIPerC) and after the ischemia (RIPostC) protect against cerebral ischemia-reperfusion injury by modulating the peripheral immunomodulation.

This study indicated that RIC neuroprotection is present in ischemic mice via the inflammatory response and metabolic changes both in the peripheral blood and ischemic brain.

## INTRODUCTION

Ischemic conditioning is a maneuver by which a series of short intervals of ischemia and reperfusion is utilized to reduce tissue damage following a prolonged ischemic event as cerebral ischemia ^1^. It protects tissue against ischemia and reperfusion injuries not only locally but also remotely from the organ being conditioned, the “so-called” remote ischemic conditioning (RIC) ^2^. At present, the way in which the peripheral signal is transmitted to a distant organ is not well understood.

Many studies have suggested that the immune and inflammatory responses in the brain and blood play critical roles in stroke pathology ^3^. Accumulating findings also indicated that remote ischemic perconditioning (RIPerC) or remote ischemic postconditioning (RIPostC) provided the neuroprotection against stroke by modulating the immune response during or after the cerebral ischemia^4, 5^. However, the effect of both strategies on the peripheral immune response has yet to be studied in detail.

In the last few years, there has been an increasing number of publications on metabolic changes related to ischemic stroke, both in patients and in animal models ^6–10^. Furthermore, animal studies of tissues or plasma have suggested ischemic conditioning to regulate metabolites involved in glycolysis and glutathione oxidation balance, among others ^11^. At this point, a full understanding of the metabolic changes and how they are involved in local as well as remote protection does, however, not exist.

We assumed that RIC may protect against cerebral injuries after stroke through potential protective mechanisms that are related with peripheral blood circulation, which are closely associated with brain inflammation. In this study, we examined the cytokine expression induced by RIPerC and RIPostC in the blood based on a multiplex cytokine assay. Furthermore, we used a non-targeted metabolomic approach to study changes in the plasma and brain metabolome of ischemic mice during the acute phase following a RIC intervention. This was expected to improve the understanding of the biology involved in the protection, as well as to identify potential novel cytoprotective metabolites.

Thus, the present study was designed to address the ischemic tolerance phenomenon by the effect of RIPerC and RIPostC on a preclinical mice model of ischemic stroke and to describe its inflammatory and metabolic signature.

## MATERIALS AND METHODS

### Mouse strain and ethical procedures

This work was conducted with the approval of the ethical committee of the University of Lleida (CEEA 02-02/20). All experiments complied with the “Principles of laboratory animal care” (NIH). Animal work is reported according to the ARRIVE guidelines ^12^.

CD1 male mice were bred in-house and maintained in the animal house facility of the University of Lleida. Young adult (2-4 months-old) mice were used and were housed in groups of two to five and separated into individual cages after surgery. They were maintained at 22±2 °C in a standard 12 h light / 12 h dark cycle. All experiments were performed during the light cycle. Food (Teklad Global 14% Protein rodent maintenance diet, Envigo, Madison, WI, USA) and water were provided *ad libitum*.

### Brain ischemia: focal transient middle cerebral artery

Animals were anesthetized with isoflurane (IsoFlo, #71002EU, Zoetis, Madrid, Spain) via facemask (5% for induction and 2% for maintenance), and body temperature was monitored by a rectal probe and maintained at 37.0±0.5°C by a thermal blanket (HB101SM451, Panlab, Cornellà, Barcelona, Spain). Mice were placed on a stereotaxic instrument (SAS-4100, ASI-Instruments, Warren, MI, USA) in prone position and eyes were protected from ocular dryness during surgery using a topical ophthalmic lubricating ointment (Lubrithal, Dechra Pharmaceuticals, Northwich, UK).

A skin incision was performed between the right eye and right ear. The temporalis muscle was dissected under the operating microscope until the temporal bone was clearly visible. After visually identifying the middle cerebral artery (MCA) under the semitranslucent skull, a hole was drilled (Ideal Micro Drill, CellPoint Scientific, Gaithersburg, MD, USA) under saline cooling. A thin layer of bone was removed carefully (1.5-2mm diameter). A flexible laser Doppler fiber (moorVMS-LDF1, Moor Instruments, Devon, United Kingdom) was placed over the skull to monitor continuously regional cerebral blood flow (rCBF). A glass capillary was used to compress the MCA for 60 min. The tip of the capillary was blunted by flame in order to avoid any damage to MCA. The compression was applied to achieve a steady decrease in rCBF below 75-80% of baseline. After 60 min of occlusion, the capillary was carefully removed to restore blood flow (reperfusion).

Animals whose rCBF did not reach to 80% of the baseline during recanalization, were excluded. The mice were allowed to recover after incision closure and were housed individually until euthanizing. All surgical procedures were performed under an operating stereomicroscope (OPMI 1-FC, Zeiss, Oberkochen, Germany).

### Experimental groups

Animals were classified into: Sham, Stroke, Stroke+RIPerC and Stroke+RIPostC groups. Sham-operated mice underwent the operation without MCA occlusion. The Stroke group underwent tMCAo by occlusion of the right MCA for 60 min, followed by reperfusion for 72h. The Stroke+RIPerC group underwent 3 cycles of ischemia for 5 min and reperfusion for 5 min after 15 min-tMCAo. The Stroke+RIPostC group underwent 3 cycles of ischemia for 5 min and reperfusion for 5 min after reperfusion for 10 min.

### Limb remote ischemic conditioning

Limb remote ischemic conditioning (LRIC) consisted of 3 cycles of right hind limb ischemia for 5 min, using an elastic gum (#430488, Corning, Glendale, AZ, USA) tightened to achieve limb pallor, followed by reperfusion for 5 min. The noninvasive LRIC procedures were performed in anesthetized mice with 2% isoflurane. Limb ischemia was confirmed by a change in the color of the skin and a decrease in limb temperature. Following limb reperfusion, the skin color returned to pink and the limb temperature to baseline.

### Multiplex cytokine analysis

Blood samples were obtained at consecutive time points: basal (before surgery), post-surgery, 6h, 24h, 48h and 72h after tMCAo surgery through submandibular puncture and collected in EDTA collection tubes (#16.444, Microvette CB300 EDTA, Sarstedt, Nümbretch, Germany). Samples were centrifugated at 10.000*xg* at room temperature for 5 min (centrifuge MiniSpin® Eppendorf, ClearLine, Venice, Italy) and stored at −80°C until further use.

Mouse High Sensitivity T Cell Magnetic Bead kits were purchased from EMD Millipore (MHSTCMAG-70KPMX, Burlington, MA, USA) and used to quantify 18 different mouse cytokines in accordance with the manufacturer’s recommendations (belonging to 4 main categories: pro-inflammatory cytokines, anti-inflammatory cytokines, chemokines and growth factors). Briefly, after pre-wetting the plates, 50 µL of each Standard or Control reagents were added into the appropriate wells. Fifty microliters of the Serum Matrix were added to the sample wells. Twenty-five microliters of Assay Buffer and 25 µL of sample (dilution 1:1) were added to the sample wells. Twenty-five microliters of pre-combined beads were added to each well. After an O/N incubation at 4°C with agitation, the plates were washed 3 times, 25 µL of Detection Antibodies was added to each well, and the plates were incubated with agitation for 1 hour at RT. Twenty-five microliters of Streptavidin-Phycoerythrin was then added to each well, and the plates were shaken for 30 min at RT. Finally, the plates were washed 3 times, and 150 µL of Drive Fluid was added to all wells. Each sample, standard, and quality control was measured in duplicate. Plates were read using a MAGPIX instrument (Luminex, Austin, TX, USA), and results were analyzed using MILLIPLEX Analyte 5.1 software (EMD Millipore, Burlington, MA, USA). For those analytes that were below the Minimum Detectable Concentration (MinDC), the lower limit of detection values divided by 2 was used as a criterion for data analysis. Cytokines contents were expressed in pg/mL.

### Non-targeted metabolomics and lipidomics analysis

#### Sample processing

Plasma samples were obtained at three different time-points: basal (before surgery), 6h and 72h after tMCAo surgery. Brain samples were obtained at 72h after tMCAo surgery and two different regions were dissected: core and contralateral. These two areas were collected into Eppendorf tubes, which were immediately flash-frozen in liquid nitrogen and stored at −80°C until further use. Both plasma and brain samples were randomized and decoded prior extraction.

Brain tissue (±25 mg of whole tissue) was homogenized using a digital ULTRA-TURRAX^®^ instrument (IKA, Staufen, Germany) in a buffer containing 180 mM KCl, 5 mM MOPS, 2 mM EDTA, 1 mM DTPAC adjusted to pH=7.4. Prior to homogenization, 1 µM BHT and a mix of proteases inhibitors (GE80-6501-23, Sigma, Madrid, Spain) and phosphatase inhibitors (1 mM Na3VO4, 1 M NaF) were added. Protein concentration was measured using the Bradford method (500-0006, BioRad Laboratories, Barcelona, Spain). Then, metabolites were extracted and lipidomic samples were prepared following the same protocol as the used for plasma samples.

#### Analytical instruments / equipment

Non-targeted metabolomic and lipidomic profiling were performed on an Agilent 1290 LC system coupled to an electrospray-ionization quadruple time of flight mass spectrometer (ESI-Q-TOF-MS/MS 6520 instrument, Agilent Technologies, Barcelona, Spain). Identities were confirmed by exact mass, retention time, isotopic distribution and/or MS/MS spectrum using public databases such as Human Metabolome Database (HMDB, http://www.hmdb.ca) and LipidMatch, a R-based tool for lipid identification^13^.

#### Non-targeted metabolomic profiling

For metabolomic analysis, plasma metabolites extraction was performed based on the methodology previously described ^14^. Briefly, 90 μl of cold methanol (containing BHT as antioxidant and phenylalanine-C13 as an internal standard) were added to 30 μl of plasma, incubated for one hour at −20°C and centrifuged at 13.000 rpm for 3 min. The supernatant was recovered, filtered with a 0.22 μm Eppendorf filter and centrifuged at 3.000 rpm for 10 min at room temperature. The resulting filtrate was transferred into vials with glass inserts (Agilent Technologies, Barcelona, Spain) for further analysis.

Two μL of extracted sample was applied onto a LC system with a reversed-phase column (Zorbax SB-Aq 2.1 x 50mm, 1.8 μm particle size, Agilent Technologies, Barcelona, Spain) equipped with a precolumn (Zorba-SB-C8 Rapid Resolution Cartridge, 2.1 × 30mm, 3.5 µm particle size, Agilent Technologies, Barcelona, Spain) and with a column temperature of 60°C. The flow rate was 0.6mL/min during 19 min. Solvent A was composed of water containing 0.2% acetic acid (v/v) and solvent B was composed of methanol 0.2% acetic acid (v/v). The gradient started at 2% of solvent B and increased to 98% B in 13 min and held for 6 min. Post-time was established in 5 min.

Data were collected using the MassHunter Data Analysis Software (Agilent Technologies, CA, USA). Quality controls (plasma samples with internal Phe-^13^C) were placed every ten samples. Peak determination and peak area integration were carried out with MassHunter Quantitative Analyses (Agilent Technologies, CA, USA).

#### Non-targeted lipidomic profiling

Briefly, to precipitate the protein fraction, 5µL of miliQ water and 20 µL of methanol were added to 10 µL of plasma sample. After the addition, samples were shaken for 2 min. Then, 250 µL of MTBE plus internal standards were added and samples were ultra-sounded in a water bath (ATU Ultrasonidos, Valencia, Spain) with a frequency and power of 40 kHz and 100 W, respectively, at 15 °C for 30 min. Then, 25 µL of miliQ water were added to the mixture and shacked for 2 min. The organic phase was separated by centrifugation (3.000 rpm) at 4°C for 10 min. The upper phase, containing all the extracted lipid’s species, was collected and subjected to mass-spectrometry. A pool of all lipid extracts was prepared and used as quality controls (QC). Ten µL of lipid extract was applied onto 1.8 µm particle 100 × 2.1 mm in Waters Acquity HSS T3 column (Waters, Mildord, MA, USA) heated at 55°C. The flow rate was 400 μl/min with solvent A composed of 10mM ammonium acetate in acetonitrile-water (40:60, v/v) and solvent B composed of 10 mM ammonium acetate in acetonitrile-isopropanol (10:90, v/v). The gradient started at 40% B and reached 100% B in 10 min and held for 2 min. Finally, the system was switched back to 40% B and equilibrated for 3 min. Electrospray ionization was performed in both positive and negative ion mode using N_2_ at a pressure of 50 psi for the nebulizer with a flow of 12L/min and a temperature of 325°C, respectively.

### Statistical analysis

For omic data, prior to statistical analyses, data was pre-treated (auto-scaled and log-transformed). Multivariate statistics was performed using Metaboanalyst software ^15^. Principal component analysis (PCA), partial least squares-discriminant analyses (PLS-DA), hierarchical clustering analysis represented by a heat map, and Random Forest used as a classification algorithm were performed using Metaboanalyst software.

Statistical analysis was performed using GrapdPad Prism (version 9.0.0) for OS X (GraphPad Software, La Jolla California, USA). Each animal was considered an independent entity. The number of mice analyzed wass indicated in the figure legend of each result. Multiple groups were compared using analysis of variance (ANOVA) test. The two-way ANOVA was used when there were two independent variables. Post-hoc test used were Dunnett’s and Tukey’s tests. Differences with *p* value <0.05 were considered significant. The specific test used in each experiment and n values are reported in the figure legends.

## RESULTS

### Plasma tMCAo related cytokines

The assessment of the cytokine profile in plasma at different periods after tMCAo showed significant changes in concentration of 14 cytokines, chemokines and growth factors such as GM-CSF, IL-1α, IL-1β, IL-4, IL-5, IL-6, IL-10, IL-12p70, IL-13, IL-17A, KC/CXCL, MCP-1, MIP-2 and TNF-α. The main source of secretion and function of each of these cytokines and chemokines is summarized in Supplementary Table 1. After ischemic stroke, cytokines were divided in two groups depending on the time in which they play a role: hyperacute phase (minutes to hours) and acute phase (hours to days) (Fig. 1).

**Figure 1:**
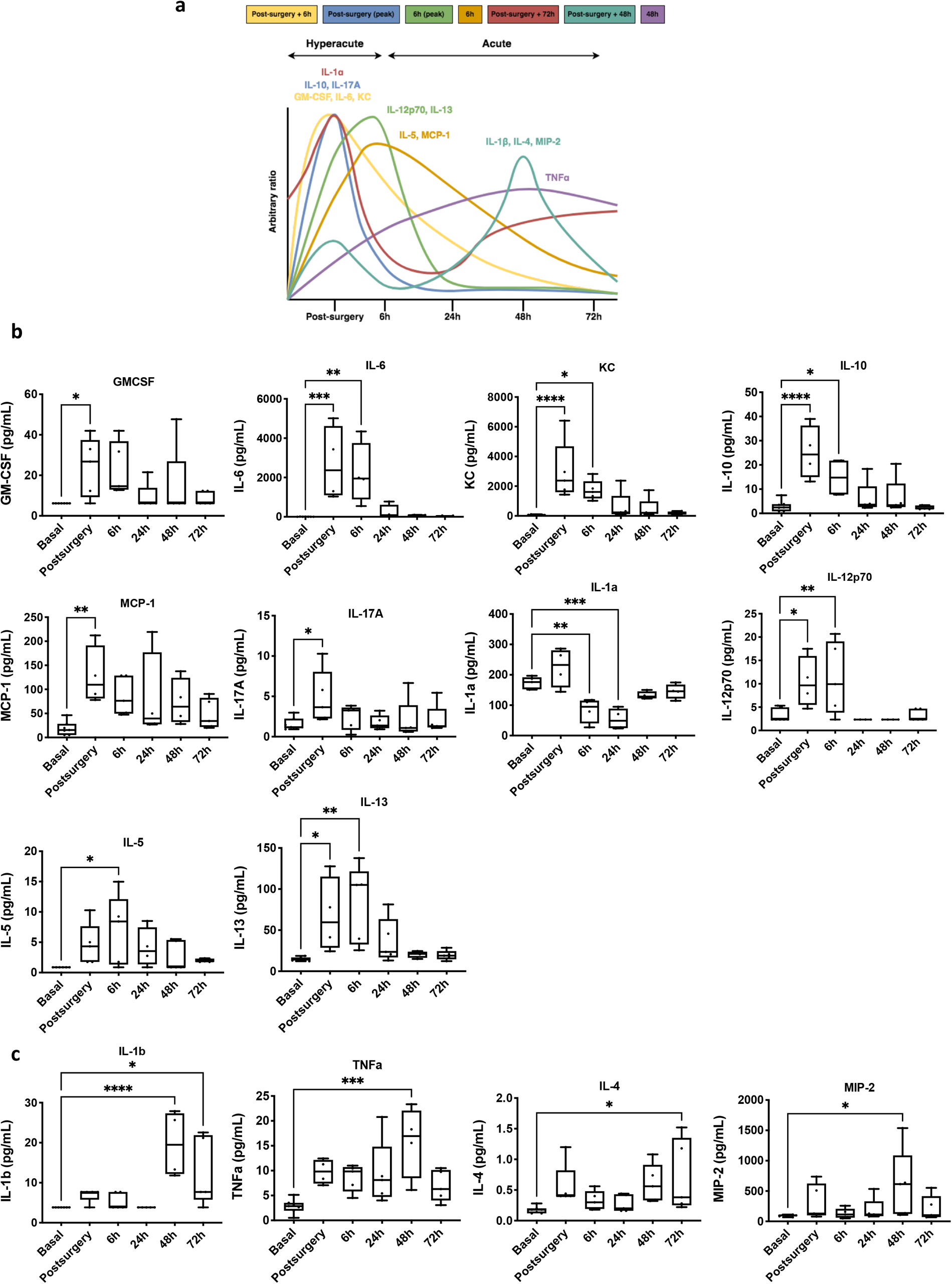
Time-depending cytokine expression profile in tMCAo mice model. **a** Time course of expression profile of cytokines, chemokines and growth factors of ischemic brain damage. Data are presented as relative increases in expression levels on an arbitrary y axis and the peak time of expression after surgery on the x axis. Cytokine, chemokine and growth factor levels in plasma samples of tMCAo mice model at hyperacute phase (**b**) and acute phase (**c**). Expression levels were analyzed at 6 different times: basal, postsurgery, 6h, 24h, 48h and 72h post-surgery. Box and whiskers graph showing min-max range, mean and all points (n=4-5 mice per group). One-way ANOVA with Dunnett’s multiple comparison test. *p<0.05, **p<0.01, ***p<0.001 and ****p<0.0001 versus Basal.

In the hyperacute phase, the expression of pro-inflammatory cytokines such as IL-6, IL-17A, IL-12p70; anti-inflammatory cytokines IL-10 and IL-13; chemokines KC and MCP-1 and growth factors GM-CSF and IL-5 increased up (Fig. 1a). In contrast, IL-1α expression levels decreased in this phase. In the acute phase, the expression of pro-inflammatory cytokines such as IL-1β and TNF-α; anti-inflammatory cytokines IL-4 and chemokine MIP-2 augmented (Fig. 1b). We did not observe significant differences in IFN-γ, LIX, IL-7 and IL-2 plasma levels.

To determine whether plasma cytokines expression defines time after stroke, multivariate statistics were applied. Non-supervised principal component analysis (PCA) suggested the existence of a different plasma cytokine profile within time, capable to explain up to 55.5% of sample’s variability (Supplementary Fig. 1a). A hierarchical clustering of the samples represented by a heat map confirmed the existence of a time-specific cytokine profile (data not shown). Furthermore, average cytokines values suggested the existence of different cytokine profile for hyperacute phase (such as postsurgery and 6h), and acute phase (24h, 48h and 72h) (Supplementary Fig. 1f). These results were confirmed by performing a supervised analysis, such as partial least squares discriminant analysis (PLS-DA) (Supplementary Fig. 1b). This allowed to determine the ability to predict the time after stroke of a specific specimen according to its cytokine profile. Cross-validation values of PLS-DA model scored a maximum Q2 of 0.38 and R2 of 0.84 when using four components (Supplementary Fig. 1c). Permutation tests (1000 repeats) yielded a low p=0.036, indicating that none of the distributions formed by the permutated data was better than the observed statistic based on the original data (Supplementary Fig. 1d). The discriminating power between groups of the different measured features was ranked by applying a variable importance projection (VIP) score. After selecting those features with VIP score >1.5 as significant, IL-4 and KC cytokines were found to be the top-ranked features (Supplementary Fig. 1e). Accordingly, RF classification algorithm revealed a time overall classification error of 0.594, being 6h, 48h and 72h the times with the highest classification error (80% of these times were not properly classified according to their cytokine plasma profile) (Supplementary Fig. 1g), being KC the cytokine with the highest contribution to classification accuracy (Supplementary Fig. 1h).

The heatmap (Supplementary Fig. 1f) confirmed the existence of a time-specific plasma cytokine profile. This suggested that there were 7 cytokine expression profiles related with stroke onset time. Figure 1c shows the time-depending cytokine expression profiles after stroke. We can clearly observe that there are two cytokine expression profiles depending on whether they are expressed in the hyperacute or acute phase of stroke. However, within these two phases, there are also subphases that are well differentiated. We classified these subtypes according to their time of expression as:

- One wave of cytokine expression at post surgery time and a progressive return to basal level [GM-CSF, IL-6 and KC] – Yellow line
- A focalized increase at post-surgery time [IL-10 and IL-17A] – Blue line
- An increase in post-surgery time and a reduction of cytokine expression at 6h up to 24h and a return to baseline level [IL-1a] – Red line
- A focalized increase at 6h time [IL-12p70 and IL-13] – Green line
- One wave of cytokine expression at 6h and a progressive return to its basal level [IL-5 and MCP-1] – Orange line
- Two waves of cytokine expression. An increase at post-surgery time and a more focalized one at 48h [IL-1β, IL-4 and MIP-2] – Aquamarine line
- A progressive increase reaching a stable expression at 24h time [TNF-α] – Violet line

### Plasma RIC-related cytokines

We further defined the inflammatory profile of RIC in the preclinical tMCAo model using the same multiplex panel of cytokines as above. The results showed that both RIC treatments had an effect mainly in the hyperacute phase after a stroke, especially in the post-surgery period (Fig. 2a). At this time point, RIPerC and RIPostC significantly decreased IL-12p70, IL-1a, IL-10, IL-13, IL-6, GM-CSF, KC and MCP1 levels compared with stroke group. At 6h, RIPerC significantly increased the levels of anti-inflammatory cytokines such as IL-10, IL-13 and GM-CSF, while decreasing KC levels compared with stroke group.

**Figure 2:**
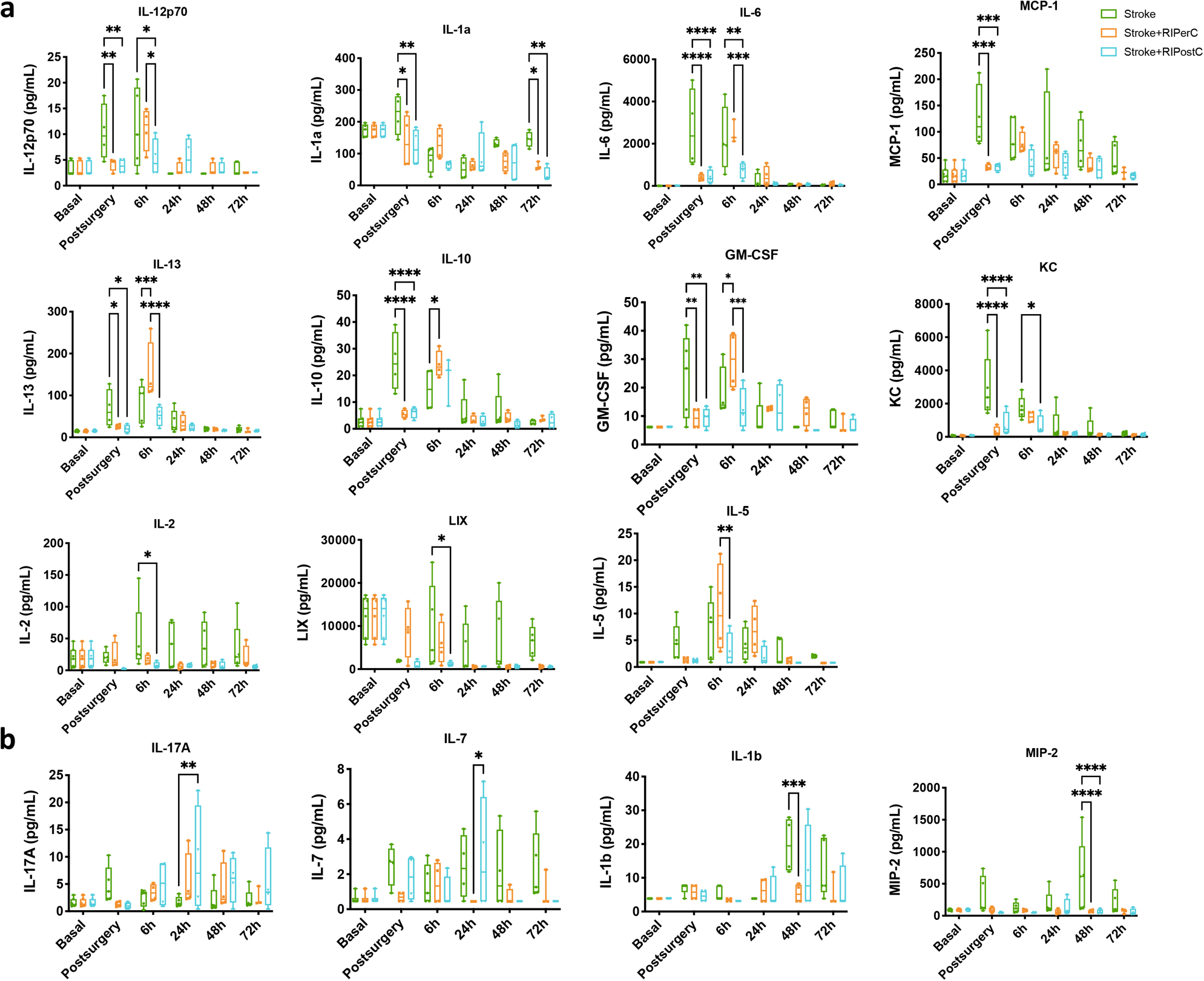
The effect of RIPerC and RIPostC on plasma levels pro-inflammatory, anti-inflammatory cytokines and growth factors in stroke animals at acute phase (**a**) and subacute phase (**b**). N=4-5 animals per group. Two-way ANOVA with Tukey’s multiple comparisons test. *p<0.05, **p<0.01, ***p<0.001 and ****p<0.0001.

At the same time, RIPostC significantly reduced the levels of IL-12p70, IL-2, IL-6 and chemokines LIX and KC. However, 6h after stroke was the time in which the greatest differences were observed between RIPerC and RIPostC. Levels of IL-13, IL-5, IL-6 and GM-CSF were significantly decreased in RIPostC group compared with RIPerC group (Fig. 2a).

Most of the differences observed at the acute phase indicated that both treatments reduced the inflammatory response. Twenty-for hours after stroke, RIPostC increased IL-17A expression compared with Stroke and IL-7 compared with RIPerC. Forty-eight hours after stroke, RIPerC reduced IL-1β and MIP-2 levels, the later cytokine was also reduced by RIPostC. Finally, both RIC treatments reduced significantly IL-1a levels at 72h after stroke (Fig. 2b). We did not observe significant differences on IFN-γ, IL-4 and TNF-α plasma levels (data not shown).

Supplementary Figure 2 represents the cytokine expression profile analyzed according to the experimental group. Taken together, these results suggested a different inflammatory signature when RIPerC and RIPostC were applied. Specifically, both treatments reduce the post-surgery inflammatory response, while at 6h both showed a specific profile. This cytokine panel has made it possible to identify plasma biomarkers, and thus to propose circulating pathophysiological mechanisms of RIC.

### Omics signatures

Next, we aimed to determine the omic (metabolome and lipidome) profile of ischemic stroke and the effect of RIPerC and RIPostC treatments in the preclinical tMCAo mouse model to describe local (brain tissue) and circulating (plasma) biomarkers. For the statistical analysis, all brain tissue databases (metabolomics and lipidomics from both ionization modes: positive and negative) were pooled (global metabolome). The same was done with plasma databases.

In brain tissue samples, our global metabolome analysis detected a total of 41.856 molecular features once baseline correction, peak picking, and peak alignment were applied on acquired data. After quality control assessment, filtering, and correcting the signal, 1.871 features remained and were used for statistical analysis. In plasma samples, our global metabolome analysis detected a total of 141.400 molecular features, of which, after processing, 2.303 features remained and were used for statistical analysis.

#### Effect of brain ischemia in the brain metabolome

Non-targeted metabolomics and lipidomics analysis were performed to investigate whether tMCAo had a brain metabolome signature. Thus, the effect of 60 min occlusion in the brain metabolome of tMCAo mice model at 72h after stroke was evaluated (Fig. 3a). Principal component analysis (PCA) showed a good separation of both groups with the three first principal components (PC1, PC2 and PC3) explaining 50% of the variability of the samples. Partial least-squares discriminant analysis (PLS-DA) was able to clearly separate the two groups (Fig. 3b), but permutation tests (1000 repeats) yielded a not significant *p* value (p=1), indicating that it is not an optimal model (data not shown).

**Figure 3:**
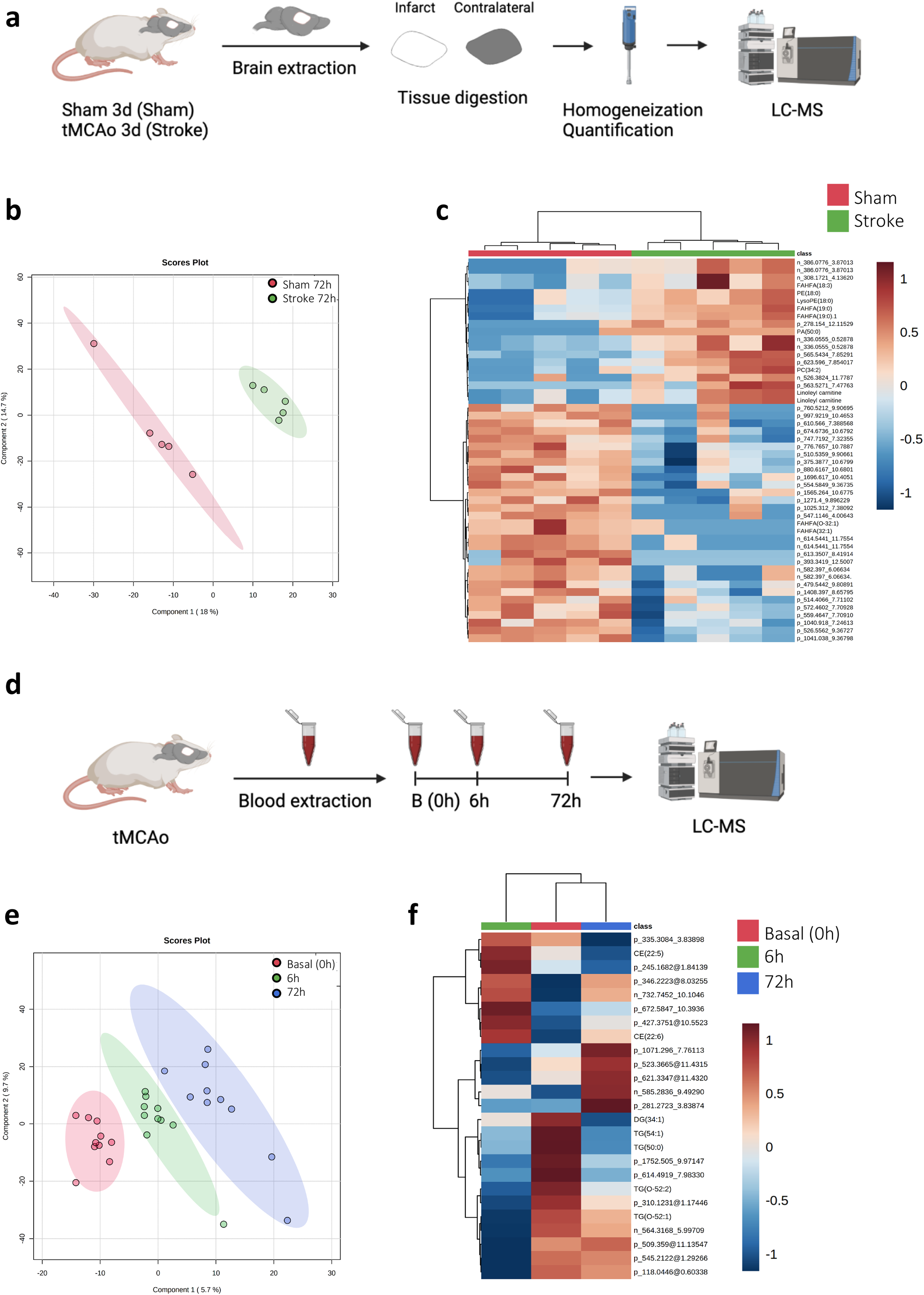
**a** Experimental design of the multivariate analysis of the differential metabolites species detected in brain tissue 72h after sham or stroke surgery (N=5 mice per group). **b** Partial-least squares discriminant analysis (PLS-DA) representation. **c** Heatmap of hierarchical clustering using the 50 metabolites with the lowest p-value detected in plasma. Each colored cell on the map corresponds to a feature and its relative concentration value, with samples in columns and compounds in rows. **d** Experimental design of the multivariate analysis of the differential metabolites detected in the plasma during basal (0h), 6h and 72h after tMCAo mice (N=10 mice per group). **e** PLS-DA representation. **f** Hierarchical clustering of average sample values.

Hierarchical clustering using all lipid and metabolites species detected showed there was no specific trend when the whole lipidome and metabolome was considered. However, when this analysis was performed using the 50 metabolites species with the lower *p* value (Fig. 3c), it confirmed that stroke affected the brain lipidome and metabolome at 72h after surgery.

A non-parametric Wilcoxon rank test was performed and from the 1871 molecules detected, 120 compounds were found to be statistically different of which 35 were identified (Table 1). Among identified species, we described 1 sterol lipid (ST), 9 fatty acids (FA), 5 glycerolipids (GL), 14 glycerophospholipids (GP), 5 sphingolipids (SP) and 1 N-acylamida. Within FAs class, 66% were FAHFAs and 34% others. Within GLs class, 40% were MGs, 40% TGs and 20% others. Within GPs class, 36% were PAs, 21% PEs, 14% LysoPCs, 14% LysoPEs and 15% others. Within SPs, 60% were ceramides and 40% others. Globally, these lipids were related with anti-oxidant, anti-inflammatory, bioenergetics, structural and signaling functions.

**Table 1:** Metabolites species significantly different in brain tissue at 72h after tMCAo between Stroke and Sham group.

| Class | Feature | m/z | RT | p.value | FDR p.value | Adduct | Sham vs Stroke<br>(Sham > Stroke = down)<br>(Sham < Stroke = up) |
| --- | --- | --- | --- | --- | --- | --- | --- |
| ST | Tauro-tetrahydroxyl bile acid | 530.2522 | 11.86807 | 0.027 | 0.74 | M-H | down |
| FA | Adrenic acid | 331.2627 | 11.872 | 0.022 | 1 | M-H | up |
| FA | FAHFA(18:3) | 307.1648 | 4.136205 | 0.016 | 0.46 | M-H | up |
| FA | FAHFA(19:0) | 327.2317 | 11.43521 | 0.003 | 0.62 | M-H | up |
| FA | FAHFA(19:0) | 327.2317 | 11.435 | 0.036 | 1 | M-H | up |
| FA | FAHFA(20:2) | 337.2083 | 13.159 | 0.016 | 0.73 | M-H | down |
| FA | FAHFA(32:1) | 507.4392 | 11.27198 | 0.01 | 0.46 | M-H | down |
| FA | FAHFA(O-32:1) | 507.4392 | 11.272 | 0.045 | 1 | M-H | down |
| FA | Linoleyl carnitine | 422.3257 | 8.4148 | 0.046 | 0.91 | M-H | up |
| FA | Linoleyl carnitine | 422.3257 | 8.414772 | 0.032 | 0.47 | M-H | up |
| GL | MG(15:0) | 297.2423 | 10.925 | 0.032 | 0.73 | M-H<sub>2</sub>O-H | down |
| GL | MG(22:0) | 432.3677 | 12.14 | 0.012 | 0.65 | M+NH<sub>4</sub> | down |
| GL | Co(Q9) | 812.6579 | 9.400449 | 0.021 | 0.55 | M+NH<sub>4</sub> | up |
| GL | TG(52:1) | 878.8198 | 10.42302 | 0.036 | 0.49 | M+NH<sub>4</sub> | up |
| GL | OxTG(62:12) | 1044.78 | 10.78937 | 0.008 | 1 | M+NH<sub>4</sub> | down |
| GP | PA(27:1) | 575.3735 | 11.67 | 0.032 | 0.73 | M-H | down |
| GP | PA(29:1) | 605.4413 | 8.005958 | 0.016 | 0.46 | M+H | down |
| GP | PA(32:1) | 647.4628 | 9.047689 | 0.008 | 0.39 | M+H | down |
| GP | PA(43:2) | 799.6193 | 6.251892 | 0.008 | 1 | M+H | down |
| GP | PA(50:0) | 901.727 | 14.615 | 0.045 | 0.89 | M+H | up |
| GP | PC(34:2) | 758.5738 | 7.053285 | 0.01 | 0.85 | M+H | up |
| GP | PE(18:0) | 480.3081 | 11.34 | 0.032 | 1 | M-H | up |
| GP | PE(40:7) | 790.545 | 6.905731 | 0.036 | 0.49 | M+H | up |
| GP | PE(P-36:4) | 724.529 | 7.241771 | 0.008 | 0.46 | M+H | up |
| GP | PS(34:2) | 740.4857 | 12.154 | 0.045 | 0.87 | M-H<sub>2</sub>O-H | up |
| GP | LysoPC(16:0) | 554.3432 | 10.84132 | 0.008 | 0.55 | M+C<sub>2</sub>H<sub>3</sub>O<sub>2</sub> | up |
| GP | LysoPC(16:0) | 554.3432 | 10.841 | 0.032 | 0.73 | M+C<sub>2</sub>H<sub>3</sub>O<sub>2</sub> | up |
| GP | LysoPE(18:0) | 480.3081 | 11.33969 | 0.002 | 0.62 | M-H | up |
| GP | OxLysoPE(20:5) | 530.2522 | 11.868 | 0.032 | 0.73 | M-H | down |
| SP | SM(27:0) | 651.5175 | 13.391 | 0.032 | 0.73 | M+HCO<sub>2</sub> | down |
| SP | SM(29:0) | 679.5492 | 13.537 | 0.036 | 0.76 | M+HCO<sub>2</sub> | up |
| SP | Cer(36:1) | 566.5506 | 7.852914 | 0.003 | 0.62 | M+H | up |
| SP | Cer(36:2) | 564.5343 | 7.477636 | 0.012 | 0.62 | M+H | up |
| SP | Hex3Cer(44:2) | 1181.882 | 13.322 | 0.016 | 0.65 | M+NH<sub>4</sub> | down |
| N-acyl | N-Eicosapentaenoyl Leucine | 415.3043 | 11.484 | 0.039 | 0.93 | M-H | down |

All compounds are putatively annotated compounds based upon physicochemical properties and/or spectral similarity with public/commercial spectral libraries. Cer: ceramides, FA: fatty acids, FAHFA: fatty acyl esters of hydroxy fatty acid, GL: glycerolipids, GP: glycerophospholipids, MG: monoacylglycerols, PA: phosphatidic acid, PC: phosphatidylcholine, PE: phosphatidylethanolamine, PS: phosphatidylserine, SM: sphingomyelins, SP: sphingolipids, ST: sterol lipids, TG: triacylglycerols,

#### Effect of brain ischemia in the plasma metabolome

The same analysis was performed to investigate the plasma metabolome signature after 60 min tMCAO (Fig. 3d). Starting with multivariate statistics, an unsupervised principal component analysis (PCA) showed that the three first principal components (PC1, PC2 and PC3) explained 27.5% of the variability of the samples, suggesting common metabolites profiles and small differences between their metabolomes. A supervised partial least-squares discriminant analysis (PLS-DA), was able to clearly separate the three groups (Fig. 3e), but permutation tests (1000 repeats) yielded a not significant *p* value (p=0.915), indicating that the PLS-DA model obtained was overfitted, probably due to the small number of variables (data not shown). Hierarchical clustering using all metabolites showed there was no specific metabolic profile for each time point when the whole metabolome was considered. However, when this analysis was performed using the 25 metabolites species with the lower p-value, it confirmed that the stroke process affected the plasma metabolomic profile as time progresses. Average metabolites abundances were plotted as a heatmap, which showed more similar profiles between basal and 72h groups compared with the 6h group (Fig. 3f). This graph also showed the profile of certain metabolites. It can be clearly seen how there were different profiles according to the abundance of these metabolites and the time of stroke. For example, among the significantly different metabolites identified, CE(22:5) increased its levels at 6h and decreased at 72h. Other examples are DG(34:1), TG(54:1) and TG(50:0) which decreased their levels at 6h and 72h gradually. In contrast, TG(O-52:1) was decreased at 6h after stroke and subsequently returns to baseline levels.

Complementary to the information above, the metabolites’ distribution within each time has been analyzed. All the molecules identified in the non-targeted analysis (86 molecules) were jointly analyzed by a Pearson correlation, and the resulting matrix was organized by hierarchical clustering (Supplementary Fig. 3a). This analysis revealed several clusters of metabolites with similar abundance patterns in time and after selecting a correlation cutoff of 0.7, four clusters have been deeply studied (Supplementary Fig. 3b). The metabolite composition of each cluster is detailed in Table 2. The first cluster, named (a) was consisted of CL, PC, SM, CE and TG. Cluster (b) was formed by 3 classes: 3-0-sulfogalactosy, OxPC and PS. Cluster (c) and (d) were made up of TG in their totality. It is important to note also that some metabolites correlated just between 2 of them. This was the case of TG(54:1) and TG(56:2), OxPC(41:4) and PS(P-39:0), TG(60:4) and CE(20:2), TG(54:6) and TG(52:4), TG(48:1) and TG(50:2), OxPC(22:4) and PC(O-20:4).

**Table 2:** Clusters analysis of circulating metabolite profiles.

| Cluster | Cluster a | Cluster b | Cluster c | Cluster d | Others |
| --- | --- | --- | --- | --- | --- |
| **Nº of species** | 13 | 6 | 3 | 3 | 12 |
| **Nº of classes** | 6 | 3 | 1 | 1 | 5 |
| **Most abundant class** | PC | OxPC | TG | TG | TG |
| **List of metabolites in each cluster** | CL(74:2) | 3-O-Sulfogalactosy | TG(58:3) | TG(O-56:3) | TG(54:1) |
|  | PC(40:5) | OxPC(40:8) | TG(51:1) | TG(O-52:1) | TG(56:2) |
|  | PC(35:1) | 3-O-Sulfogalactosy | TG(O-54:2) | TG(O-50:0) | OxPC(41:4) |
|  | SM(34:2) | OxPC(34:2) |  |  | PS(P-39:0) |
|  | PC(32:1) | OxPC(38:7) |  |  | TG(60:4) |
|  | CL(81:2) | PS(40:3) |  |  | CE(20:2) |
|  | CL(78:2) |  |  |  | TG(54:6) |
|  | PC(34:1) |  |  |  | TG(52:4) |
|  | SM(40:1) |  |  |  | TG(48:1) |
|  | CE(22:4) |  |  |  | TG(50:2) |
|  | PC(36:4) |  |  |  | OxPC(22:4) |
|  | PC(37:2) |  |  |  | PC(O-20:4) |
|  | TG(50:3) |  |  |  |  |

Univariant metabolomic analysis demonstrated the existence of specific changes in plasma metabolome after stroke. In order to identify those entities responsible for a specific metabolic profile for each time, a non-parametric Kruskal Wallis test was performed followed by Dunn’s multiple comparison post hoc test to compare all the groups between them. Thus, when we searched for specific stroke biomarkers in plasma, we found 223 compounds to be statistically different between groups (p<0.05) (Table 3). Among all the significant molecules, 125 were annotated, and 98 were not identified. Of all identified species, we detected 8 FA, 54 GL, 41 GP, 10 SP, 7 ST, 1 benzoid, 1 amino acid (AA), 1 imidazole, 1 acylamida and 1 purine. Within the FAs class, 50% of features were acylcarnitines and the rest of the features were grouped together as others. Within GLs class, 80% of features were TGs, 13% DGs and 7% MGs. Within GPs class, 12% were cardiolipins, 32% LysoPCs, 34% PCs and 22% others. Within SPs class, 50% were sphingomyelins, 20% ceramides, 20% gangliosides and 10% others. Within STs class, 71% were cholesteryl esters and 29% others. Globally, these metabolites were related with anti-oxidant, anti-inflammatory, bioenergetics, structural and signaling functions. Metabolic alteration by tMCAo was primarily associated with FA (AcCa), GL (DG, MG and TG), GP (LysoPC, CL, PC and OxPC), SP (SM) and ST (CE) metabolic pathways. AcCa and cholesterol ester metabolism was changed most significantly at 6h. The levels of TGs decreased at 6h and 72h. The levels of DG, MG and LysoPC decreased at 6h, but increased again to basal levels by 72h. Long CL, PC and OxPC reduced at 72h. Finally, short CL as well as SM reduced at 6h but increased by 72h.

**Table 3:**
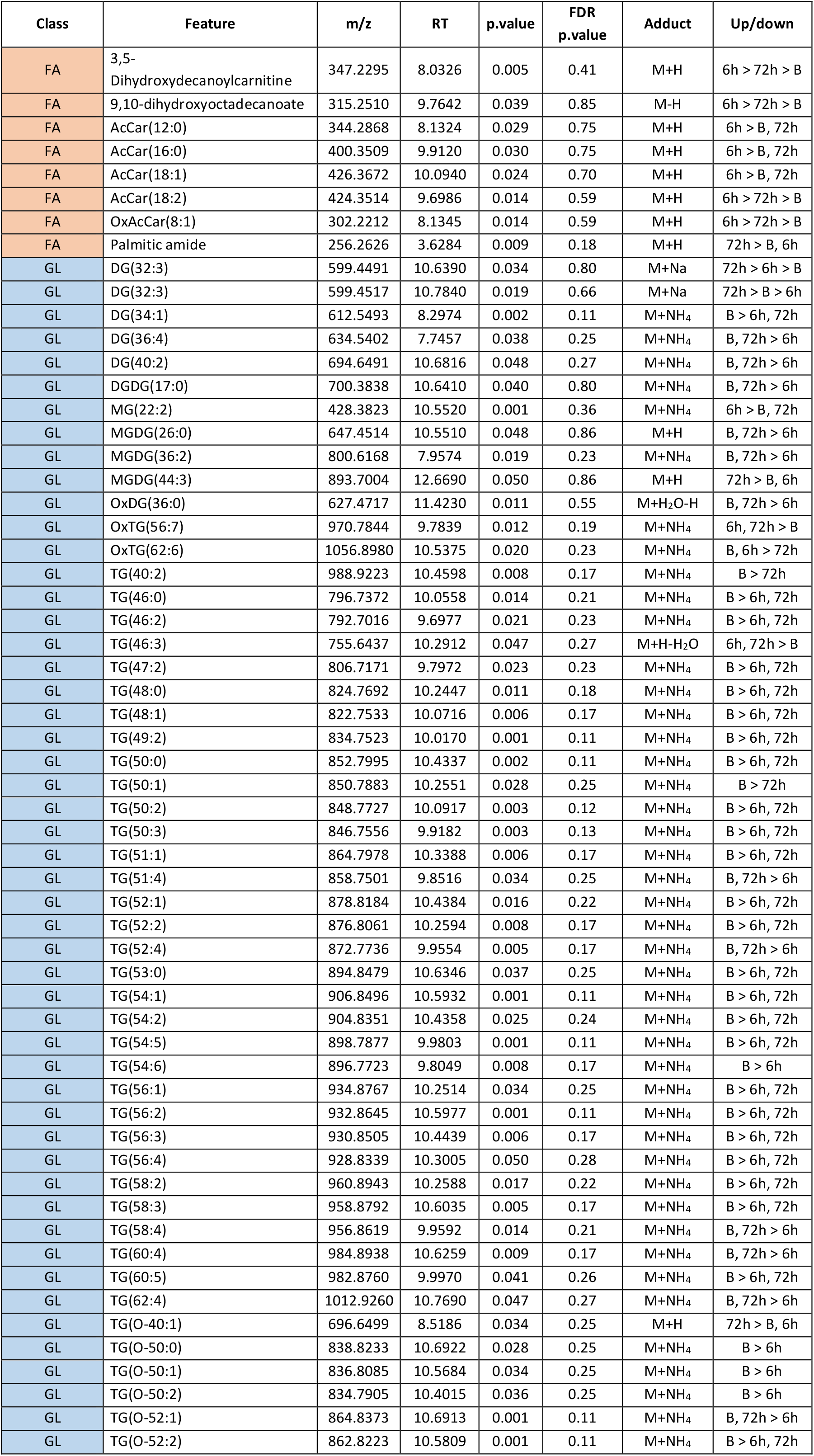
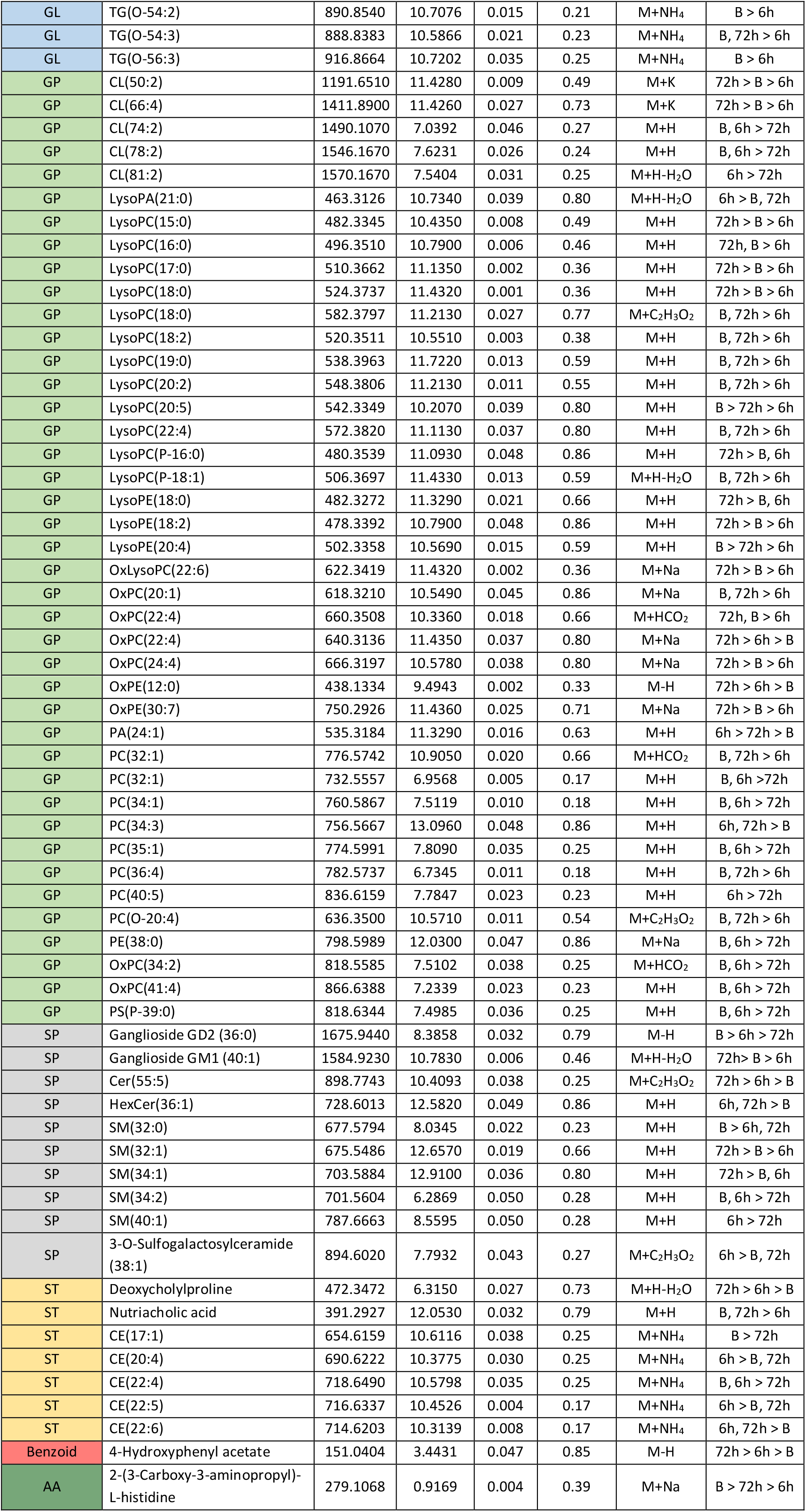
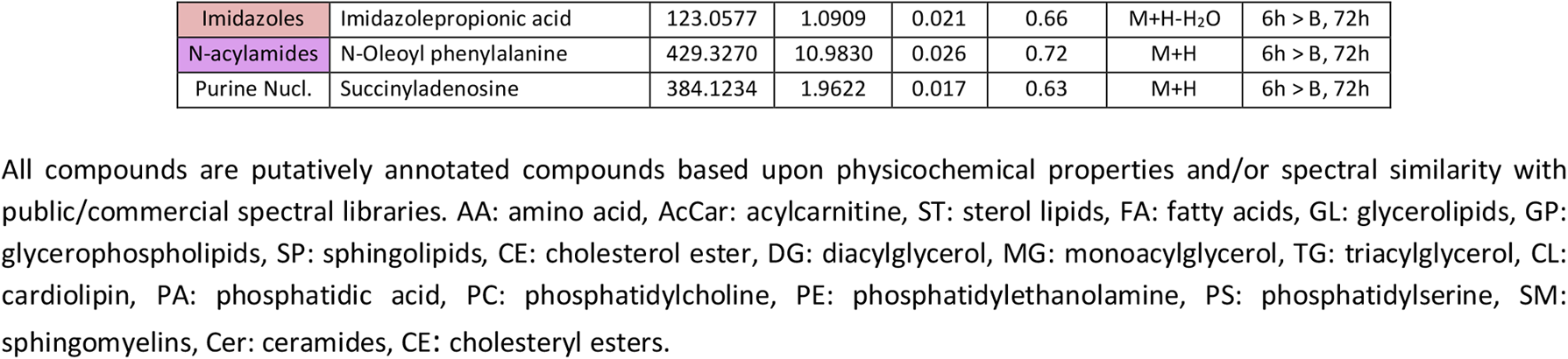
Metabolites species significantly different in plasma between basal (B), 6h and 72h after stroke.

#### Effect of RIC in brain metabolome

To investigate whether RIPerC and RIPostC had a brain metabolome signature expressing an adaptive metabolic response with a potential neuroprotective role, the same analysis was performed (Fig. 4a). Principal component analysis (PCA) showed that the three first components (PC1, PC2 and PC3) explained 36.4% of the variability of the samples. Partial least-squares discriminant analysis (PLS-DA) was able to clearly separate the three groups (Fig. 4b), but permutation tests (1000 repeats) yielded a not significant *p* value (p=0.976), indicating that the model is overfitted (data not shown). Hierarchical clustering using all lipid and metabolites species detected showed there was no specific trend when the whole lipidome and metabolome was considered. However, when this analysis was performed using the 25 metabolites species with the lower *p* value, it showed that both RIC groups (RIPerC and RIPostC) were more homogeneous and similar to each other that to the Stroke group, indicating that RIC is determinant in the definition of the metabolome in the brain (Fig. 4c).

**Figure 4:**
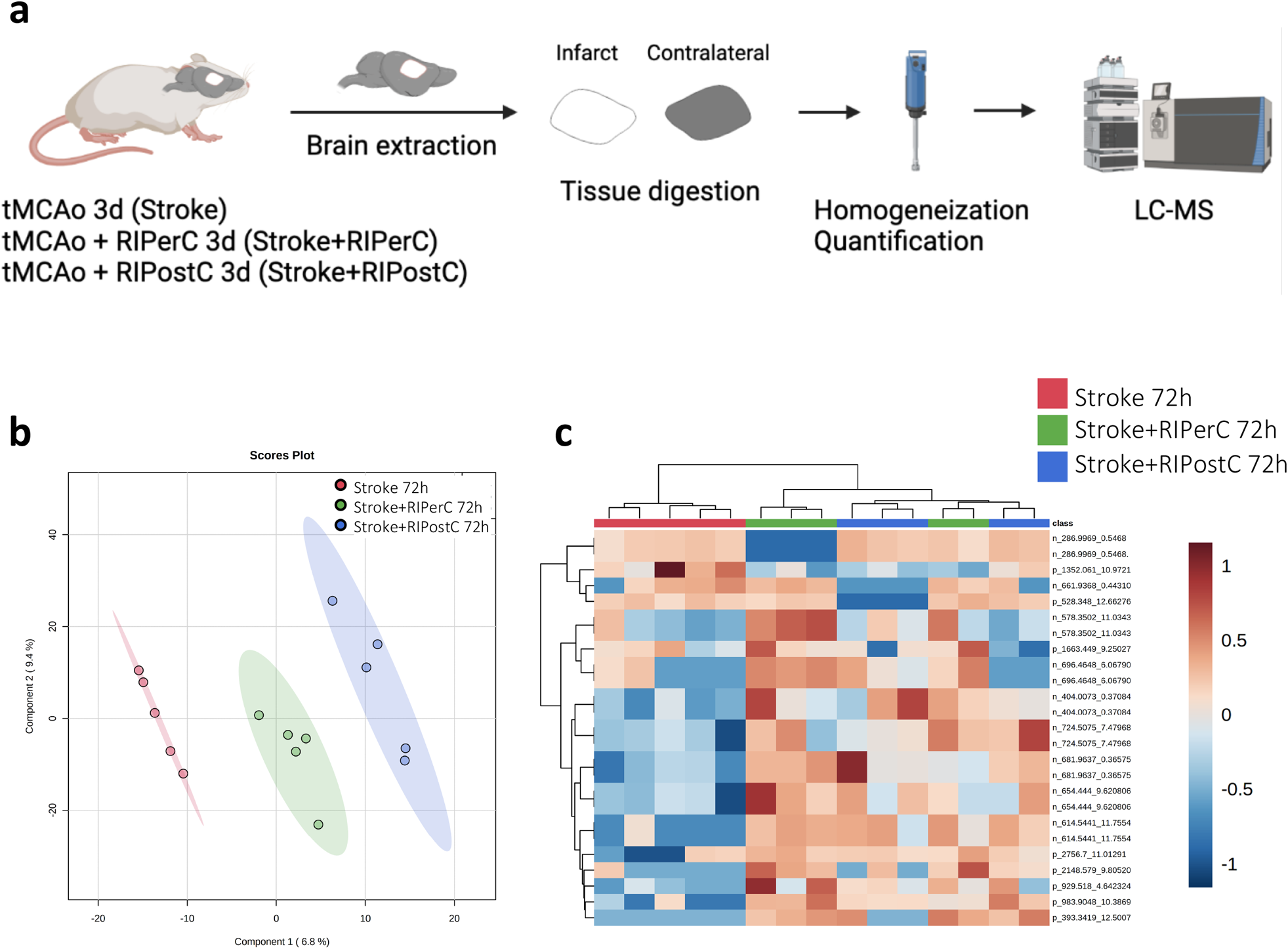
**a** Experimental design of the multivariate analysis of the differential metabolites detected in brain tissue 72h after RIC application (N=5 mice per group). **b** PLS-DA representation. **c** Heatmap of hierarchical clustering using the 25 metabolites with the lowest p-value detected in plasma.

A non-parametric Kruskal Wallis test was performed followed by Dunn’s multiple comparison post hoc test to compare all the groups. From the 1871 molecules detected, 10 compounds were found to be statistically different, of which 3 were identified (Table 4). Among identified species, we described 1 glycerolipid (GL) and 2 sphingolipids (SP). Cer(42:3) and HexCer(36:1) levels increased at RIPostC group compared to that in the Stroke and RIPerC groups. TG(28:0) levels increased at RIPerC group compared to that in the Stroke and RIPostC groups. Globally, these lipids are related with signaling, structural and bioenergetics functions.

**Table 4:** Metabolites species significantly different in brain tissue at 72h after tMCAo between Stroke, Stroke+RIPerC and Stroke+RIPostC groups.

| Class | Feature | m/z | RT | p.value | FDR p.value | Adduct | Up/down |
| --- | --- | --- | --- | --- | --- | --- | --- |
| SP | Cer(42:3) | 684.5643 | 12.5930 | 0.032 | 0.51 | M+K | Post > Stk, Per |
| SP | HexCer(36:1) | 728.5888 | 12.6010 | 0.035 | 0.89 | M+H | Post > Stk, Per |
| GL | TG(28:0) | 544.4576 | 8.8998 | 0.030 | 0.96 | M+NH<sub>4</sub> | Per > Stk, Post |

All compounds are putatively annotated compounds based upon physicochemical properties and/or spectral similarity with public/commercial spectral libraries. SP: sphingolipids, GL: glycerolipids, Cer: ceramides, TG: triacylglycerids.

#### Effect of RIC in plasma metabolome

Finally, we evaluated the effect of RIC in the plasma metabolome of tMCAo mice model at 6h (Fig. 5a) and 72h after stroke (Fig. 5d). If we focused first at 6h, principal component analysis (PCA) showed that the three first components (PC1, PC2 and PC3) explained 24.3% of the variability of the samples. Partial least-squares discriminant analysis (PLS-DA) was able to clearly separate the three groups (Fig. 5b), but permutation tests (1000 repeats) yielded a not significant *p* value (p=1), indicating that the model is overfitted (data not shown). Hierarchical clustering using all metabolites species detected showed there were no global changes when the whole metabolome was considered. However, when this analysis was performed using the 25 metabolites species with the lowest *p* value, it revealed that there was an almost perfect clusterization according to their group. The heatmap of metabolites abundances showed that the Stroke+RIPerC group had a specific profile compared with the other two groups (Stroke and Stroke+RIPostC) (Fig. 5c).

**Figure 5:**
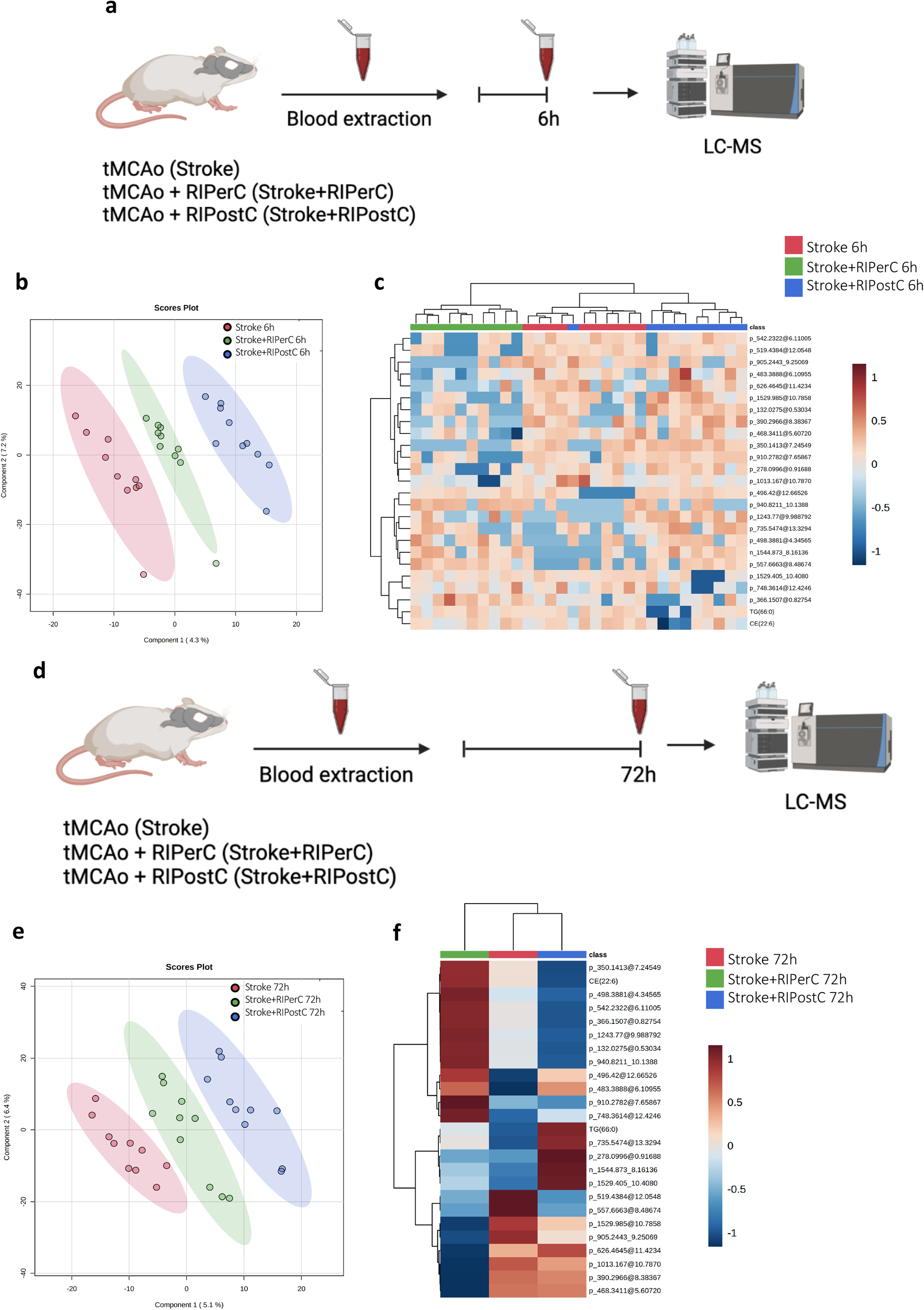
**a** Experimental design of the multivariate analysis of the differential metabolites species detected in the plasma after RIC application in the tMCAo mice at 6h after stroke (N=10 mice per group). **b** PLS-DA representation. **c** Heatmap of hierarchical clustering using the 25 metabolites with the lower p-value detected in plasma. d Experimental design of the multivariate analysis of the differential metabolites species detected in the plasma after RIC application in the tMCAo mice at 72h after stroke (N=10 mice per group). b PLS-DA representation. c Hierarchical clustering of average sample values.

Univariant metabolomic analysis demonstrated the existence of specific changes in plasma metabolome at 6h after RIC application. Thus, to search for specific RIC biomarkers in plasma at the hyperacute phase, a non-parametric Kruskal Wallis test was performed followed by Dunn’s multiple comparison post hoc test to compare all the groups. We found 61 compounds to be statistically different between groups (p<0.05). Among all the significant molecules, 18 were annotated, and 43 were not identified (Table 5). Of all identified species, we described 1 amino acid (AA), 4 glycerolipids (GL), 9 glycerophospholipids (GP), 2 N-acylamidas, 1 organonitrogen and 1 sphingolipid (SP). Within GLs class, 50% were MGs and 50% DGs. Within GPs class, 22% were PGs, 22% PEs and 56% others. In general, lipids detected were associated with anti-oxidant, signaling and structural functions.

**Table 5:** Metabolites species significantly different at 6h after tMCAo between Stroke (Stk), Stroke+RIPerC (Per) and Stroke+RIPostC (Post).

| Class | Feature | m/z | RT | p.value | FDR<br>p.value | Adduct | Up/down |
| --- | --- | --- | --- | --- | --- | --- | --- |
| AA | Oxidized glutathione | 613.1720 | 0.6303 | 0.045 | 1 | M+H | Per > Post > Stk |
| GL | MGDG(29:5) | 679.4733 | 11.0300 | 0.026 | 1 | M+H | Per > Stk, Post |
| GL | MGDG(40:0) | 843.7178 | 14.4390 | 0.030 | 1 | M+H | Stk, Post > Per |
| GL | DG(34:0) | 614.5677 | 8.6497 | 0.033 | 1 | M+NH<sub>4</sub> | Post > Stk, Per |
| GL | OxDG(32:5) | 598.3924 | 10.9160 | 0.045 | 1 | M+Na | Per > Stk, Post |
| GP | CDP-DG(50:0) | 1244.7770 | 9.9888 | 0.017 | 1 | M+K | Post > Per > Stk |
| GP | LysoPE(18:0) | 482.3683 | 11.2020 | 0.005 | 1 | M+H | Post > Per > Stk |
| GP | LysoSM(18:0) | 467.3692 | 6.6493 | 0.013 | 1 | M+H | Post > Stk > Per |
| GP | PA(21:0) | 495.2795 | 12.0470 | 0.041 | 1 | M+H | Per > Stk > Post |
| GP | PE(39:3) | 806.5215 | 12.6160 | 0.032 | 1 | M+H | Stk, Per > Post |
| GP | PE(P-42:5) | 806.6228 | 8.9158 | 0.037 | 1 | M+H | Per > Post |
| GP | PG(32:2) | 736.5546 | 13.3290 | 0.011 | 1 | M+NH<sub>4</sub> | Post > Per > Stk |
| GP | PG(36:0) | 723.5009 | 11.0680 | 0.013 | 1 | M+H | Per > Stk > Post |
| GP | PS(44:4) | 878.6416 | 11.5170 | 0.047 | 1 | M+H<sub>2</sub>O-H | Post > Per > Stk |
| N-acylamida | N-Eicosapentaenoyl Glutamine | 413.2755 | 12.0680 | 0.021 | 1 | M+H-H<sub>2</sub>O | Per > Stk > Post |
| N-acylamida | N-Linoleoyl Glutamine | 391.3038 | 8.3837 | 0.020 | 1 | M+H-H<sub>2</sub>O | Post > Stk > Per |
| Organonitrogen | Choline | 104.1073 | 11.4350 | 0.045 | 1 | M+H | Post > Per > Stk |
| SP | Ganglioside GM1 (34:0) | 1499.8380 | 8.0783 | 0.021 | 0.99 | M-H<sub>2</sub>O-H | Per, Post > Stk |

All compounds are putatively annotated compounds based upon physicochemical properties and/or spectral similarity with public/commercial spectral libraries. AA: amino acids, GL: glycerolipids, GP: glycerophospholipids, SP: sphingolipids, MG: monoacylglycerids, DG: diacylglycerids, PG: phosphatidylglycerol, PA: phosphatidic acid, PE: phosphatidylethanolamine, PS: phosphatidylserine.

Glycerophospholipid metabolism was changed most significantly in the RIPostC group compared to that in the RIPerC and Stroke group. The levels of AA and MGDG increased at RIPerC group. Sphingolipid metabolism was changed in both RIC groups compared to that in the Stroke group.

Considering the results at 72h, principal component analysis (PCA) showed that the three first components (PC1, PC2 and PC3) explained 27% of the variability of the samples. Partial least-squares discriminant analysis (PLS-DA) was able to clearly separate the three groups (Fig. 5e), but permutation tests (1000 repeats) yielded a not significant *p* value (p=0.915), indicating that, globally, metabolome was not affected by the treatment (data not shown). Hierarchical clustering using all metabolites species detected showed there was no affectation of the metabolome at a global level. Moreover, when this analysis was performed using the 25 metabolites species with the lower *p* value, the distribution was not improved. However, the heatmap of average 25 metabolites abundances allowed to see how molecules evolve depending on the treatment, with their respective abundances. A specific profile for each group can be observed (Fig. 5f).

Univariate metabolomics analysis demonstrated the existence of specific changes in the global plasma metabolome at 72h after RIC application. Thus, when we searched for specific stroke biomarkers in plasma, a non-parametric Kruskal Wallis test was performed followed by Dunn’s multiple comparison post hoc test to compare all the groups. We found 66 compounds to be statistically different between groups (p<0.05) (Table 6). Among all the significant molecules, 29 were annotated, and 37 were not identified. Of all identified species, we described 1 amino acid (AA), 1 benzoid, 3 fatty acids (FA), 9 glycerolipids (GL), 10 glycerophospholipids (GP), 3 sphingolipids (SP), 1 N-acylamida and 1 purine. Within GLs class, 78% were TGs, 11% MGs and 11% DGs.

**Table 6:** Metabolites species significantly different at 72h after tMCAo between Stroke, Stroke+RIPerC and Stroke+RIPostC.

| Class | Feature | m/z | RT | p.value | FDR<br>p.value | Adduct | Up/down |
| --- | --- | --- | --- | --- | --- | --- | --- |
| AA | L-Tryptophan | 205.0969 | 1.2644 | 0.046 | 1 | M+H | Stk, Per > Post |
| Benzoids | 2,4-Dihydroxybenzophenon | 237.0484 | 3.4239 | 0.004 | 1 | M+Na | Stk > Per, Post |
| FA | Butyryl-CoA | 860.1612 | 9.1386 | 0.022 | 1 | M+Na | Stk > Per > Post |
| FA | OxAcCa(18:0) | 444.3774 | 10.016 | 0.014 | 1 | M+H | Stk > Per > Post |
| FA | OxFA(18:0) | 283.2689 | 11.774 | 0.024 | 1 | M+H-H<sub>2</sub>O | Per > Stk > Post |
| GL | DG(34:2) | 629.5062 | 12.639 | 0.023 | 1 | M+Na | Post > Stk, Per |
| GL | MGDG(43:3) | 879.6841 | 12.666 | 0.031 | 1 | M+H | Post > Per > Stk |
| GL | TG(44:0) | 768.7036 | 9.832797 | 0.041 | 0.85 | M+NH<sub>4</sub> | Stk, Per > Post |
| GL | TG(51:3) | 860.7688 | 10.02831 | 0.009 | 0.85 | M+NH<sub>4</sub> | Per, Post > Stk |
| GL | TG(56:7) | 922.7838 | 9.920932 | 0.03 | 0.85 | M+NH<sub>4</sub> | Per > Stk, Post |
| GL | TG(O-52:0) | 887.7785 | 9.883188 | 0.027 | 0.85 | M+K | Per > Stk > Post |
| GL | OxTG(54:5) | 946.7843 | 9.840236 | 0.005 | 0.85 | M+NH<sub>4</sub> | Stk, Per > Post |
| GL | OxTG(56:5) | 974.8131 | 10.00853 | 0.032 | 0.85 | M+NH<sub>4</sub> | Per > Stk, Post |
| GL | OxTG(58:1) | 1010.905 | 10.19607 | 0.019 | 0.85 | M+NH<sub>4</sub> | Post > Stk, Per |
| GP | CL(46:0) | 1123.69 | 10.345 | 0.043 | 1 | M+Na | Post > Stk, Per |
| GP | PC(P-36:5) | 764.5779 | 7.976064 | 0.02 | 0.85 | M+H | Per > Stk |
| GP | OxPC(38:7) | 864.5413 | 6.863933 | 0.017 | 0.85 | M+HCO<sub>2</sub> | Per > Post |
| GP | OxPC(40:8) | 890.5625 | 6.9239575 | 0.025 | 0.85 | M+HCO<sub>2</sub> | Stk, Per > Post |
| GP | OxLysoPC(15:1) | 512.3456 | 9.2557 | 0.022 | 1 | M+H | Stk > Post > Per |
| GP | OxLysoPC(18:1) | 586.2908 | 9.4929 | 0.049 | 1 | M+H | Stk > Per > Post |
| GP | OxLysoPC(20:2) | 580.3596 | 10.347 | 0.041 | 1 | M+H | Post > Stk, Per |
| GP | OxLysoPC(20:4) | 608.336 | 11.141 | 0.034 | 1 | M+H | Per > Post > Stk |
| GP | PA(32:1) | 647.4648 | 9.079067 | 0.037 | 0.85 | M+H | Per, Post > Stk |
| GP | PS(40:3) | 840.55 | 6.9992015 | 0.047 | 0.87 | M-H | Per > Stk > Post |
| SP | Ganglioside GM3(30:0) | 1099.691 | 10.349 | 0.042 | 1 | M+H | Stk > Per, Post |
| SP | SM(42:9) | 799.5427 | 12.734 | 0.013 | 1 | M+H | Stk > Per > Post |
| SP | 3-O-Sulfogalactosylceramide (36:1) | 866.5555 | 7.2220205 | 0.016 | 0.85 | M+C<sub>2</sub>H<sub>3</sub>O<sub>2</sub> | Stk, Per > Post |
| N-acylamida | N-Eicosapentaenoyl Lysine | 469.2947 | 12.114 | 0.013 | 1 | M+K | Per > Stk > Post |
| Purines | Hypoxanthine | 137.0459 | 0.4506 | 0.046 | 1 | M+H | Per > Stk > Post |

All compounds are putatively annotated compounds based upon physicochemical properties and/or spectral similarity with public/commercial spectral libraries. AA: amino acids, FA: fatty acids, GL: glycerolipids, GP: glycerophospholipids, SP: sphingolipids, TG: triacylglycerids, MG: monoacylglycerids, CL: cardiolipins, PC: phosphatidylcholine, PA: phosphatidic acid, PS: phosphatidylserine, SM: sphingomyelin.

Within GPs class, 30% were PCs, 40% were OxLysoPCs and 30% others. Glycerolipids and glycerophospholipids were the most abundant class with functions such as bioenergetics and signaling. FA metabolism was most significantly changed in the RIPostC group, while GL and GP metabolism were most altered in the RIPerC group.

## DISCUSSION

We previously demonstrated that ischemic animals with RIPerC and RIPostC had a reduction in infarct volume and better neurological outcome in comparison with ischemic animals without RIC ^16^. Interestingly, our study provides for the first-time evidence of an inflammatory and metabolomic-lipidomic signature of RIC. This signature suggests that RIPerC and RIPostC protect against cerebral ischemia-reperfusion injury through the peripheral immunomodulation and by the regulation of lipid metabolism.

In this study, we showed a rapid and transient peripheral inflammatory response dominated by several inflammatory mediators after experimental stroke. These inflammatory factors circulate to the periphery and become the “invisible hand” that connects the center to the periphery after an ischemic stroke. Regarding which cytokines and chemokines were increased in mice in hyperacute stroke (until 6h), we found GM-CSF, IL-1α, IL-5, IL-6, IL-10, IL-12p70, IL-13, IL-17A, KC/CXCL1 and MCP-1/CCL-2. On acute phase (from 24h to 72h), we found that there was a substantial resolution of the inflammatory response to stroke. However, there were still several cytokine levels increased, such as IL-1ß, IL-4, MIP-2/CXCL2 and TNF-α.

In view of this complex inflammatory environment, we defined the effect of RIC in the preclinical tMCAo model. In our results, we observed differences between RIC groups and tMCAo group, concretely at postsurgery time. At this time, we observed a decrease in several inflammatory mediators in both RIC groups. These results suggest that RIC may transfer a protective signal to the ischemic brain via the peripheral immune system. We therefore hypothesize that with less chemokine expression (GM-CSF, KC and MCP-1), there is less leukocyte infiltration and in turn less pro-inflammatory response (IL-12p70, IL-1α and IL-6). This would explain also why there is also a decrease in anti-inflammatory cytokines expression (IL-10 and IL-13).

However, in later time-points we showed different expressions patterns of RIPerC and RIPostC. For instance, 6 hours after tMCAo, RIPerC significantly increase the levels of anti-inflammatory cytokines (IL-10 and IL-13), like as GM-CSF. A possible explanation for these observations is that after ischemic injury, GM-CSF chemokine expression increases to initiate leukocyte migration, but when RIPerC is applied, there is an increased anti-inflammatory response to counteract the proinflammatory response.

At the same time, RIPostC significantly reduced the levels of other pro-inflammatory cytokines (IL-12p70, IL-2 and IL-6) and chemokines (LIX and KC). This is consistent with a previous study, in which it was reported that RIPostC decreased the expression of some of the mentioned cytokines after stroke ^5^. However, these observations suggest that the effect is similar, a decrease in leukocyte infiltration and lesser pro-inflammatory response, but both treatments have different signaling pathways.

An important aspect to highlight is that 24h after ischemia, RIPostC increased plasma IL-17A expression levels. This interleukin is a solid link between innate and adaptive immunity and can exert both deleterious and beneficial effects in neuroinflammation ^17^. Thus, Lin and colleagues^17^ showed that IL-17A from reactive astrocytes maintained and augmented the survival and neuronal differentiation of neural precursor cells in the SVZ and subsequent synaptogenesis and spontaneous recovery. Therefore, although IL-17A is well known for its damage role in acute stroke, it may be an essential mediator for ischemia-induced neurorepair and spontaneous recovery after stroke.

Forty-eight hours after tMCAo, RIPerC reduced IL-1ß expression levels, evidencing that there is likely to be less neutrophil chemoattraction and therefore less tissue damage. Also, at this time point, both treatments decrease MIP-2 expression levels, suggesting that there is less recruitment of immune cells. Finally, 72h after stroke, RIPerC and RIPostC significantly reduce IL-1α indicating that there is less pro-inflammatory response and neutrophil infiltration, as well as less BBB damage.

In our study, a comprehensive metabolomic analysis of mouse brain and plasma was performed to learn the tMCAo-related changes of metabolites in the brain and periphery, as well as to assess the effect of RIC intervention in order to evaluate whether the effect verified in the reduction of infarct can be explained at a metabolomic level. Globally, the plasma seems to be more affected by the ischemic process than the brain itself suggesting a systemic reactive response.

We described a brain lipid profile of mice who had suffered ischemic stroke characterized by a significant increase in selected FA, GP and SP, mainly involved in anti-inflammatory, remodeling and signaling roles. FAs accumulation in the brain has been associated with several neurodegenerative disorders ^18^. Alterations in brain lipids develop immediately upon ischemic injury through fatty acid release by the phospholipase A2 family (PLA_2_). Adrenic acid is the third most important PUFA in the brain, after DHA and ARA. Membrane phospholipids are converted to adrenic acid, indicating that this pathway is promoted. Indeed, in the brain, linoleyl carnitine was elevated 72h after tMCAo. It is well known that ischemic conditions inhibit β-oxidation of fatty acids and leads to accumulation of toxic intermediates of β-oxidation, particularly long-chain acylcarnitine compounds like linoleyl carnitine ^19^.

GPs are the main components of neural (neurons and glia) cell membranes. Specially, PA is the precursor for “de novo” generation of different membrane GP such as PC, PE and PS. In the brain, PA species were downregulated compared to other species, suggesting its metabolic contribution to membrane biogenesis. Another GP that was upregulated in the brain 72h after tMCAo was LysoPC(16:0). It has been demonstrated that this metabolite is neuroprotective in brain ischemia models ^20^. It is also known that is a powerful inducer of superoxide ion production in endothelial cells, and that controlled levels of superoxide function as angiogenic factors in ischemic angiogenesis. Due to its neuroprotective properties, increased brain levels of LysoPC(16:0) could contribute to angiogenesis after tMCAo.

Ceramides are a type of sphingolipids whose levels increased after tMCAo. Our results agree with previous studies showing that in mice subjected to 60 minutes of tMCAo, ceramide levels were significantly increased in the ischemic cerebral cortex in acute phase ^21^. These findings establish an essential role for the ceramide pathway in the exacerbation of ischemic injury.

A bile acid that was found to be reduced in the brain was tauro-tetrahydroxyl bile acid. Bile acids attenuate neuroinflammation through neuronal apoptosis inhibition and microglia functioning. This reduction may indicate that 72h after tMCAo the inflammatory response within the brain is still present.

Acylcarnitines transport fatty acids into mitochondria for β-oxidation and energy metabolism ^22^. During β-oxidation, L-carnitine is one of the key metabolites that transports cytosolic fatty acids across the inner mitochondrial membrane, by forming acylcarnitines. Therefore, elevated circulatory acylcarnitines may reflect dysregulation of mitochondrial β-oxidation and activation of proinflammatory signaling. The increase of plasma acylcarnitines 6h after tMCAo probably reflects an increased energy requirement due to negative feedback from the brain hypoxic state. These results demonstrate that there is an increase in fatty acid catabolism in the acute phase after tMCAo to cover energy requirement.

Cholesterol is the most abundant member of the sterol lipid category in mammals. Besides cholesterol, the category is comprised, among others, by cholesterol esters. Esterification of cholesterol is a universal mechanism to store and transport large quantities of cholesterol between organs and tissues and to reduce the toxicity of the excess cellular cholesterol ^23^. Seventy-two hours after ischemia, there was a reduction of several cholesterol esters in plasma, indicating that there has probably been massive transport of these molecules to the brain to repair cell membranes damaged by ischemia.

TGs are the main constituent of animal fats and play an important role in metabolism as energy sources and transporters of dietary fats. They accumulate inside an intracellular organelle, called lipid droplets. We can observe that there is a considerable decrease at 6h of many TGs, which is mostly maintained until 72h. Little has been reported to date on the role of TGs in acute stroke and their role in poststroke recovery. A previous study demonstrated that lower TGs levels seem to be associated with a worse prognosis in AIS ^24^. One hypothesis for our results may be that low TGs in plasma are a surrogate marker of poor outcome after tMCAo. Regarding DGs, they are very important molecules as they are building blocks for GPs. Low levels of DGs were seen 6h after tMCAo, suggesting that circulating DGs in the plasma are required in the brain to recover damaged cell membranes.

In addition, 6h after tMCAo, there was a reduction of several LysoPC. One recent study have discovered that lower LysoPC plasma levels are linked to poor disease outcomes. Decreased levels of LysoPC were observed in diabetes, Alzheimer disease, and aging, and were associated with increased mortality risk ^25^. The mechanism responsible for the reduction in circulating LysoPC is unknown but may be due to a direct degradation of circulating LysoPC or enhanced clearance from the circulation by metabolically active tissues. Another explication could be that, after an ischemic event, plasma LysoPC is accumulated in the brain to produce acetylcholine. Our group previously demonstrated that low concentrations of a specific LysoPC [LysoPC(16:0)] were significantly associated with stroke recurrence ^7^. In the present study, we also observed low levels of this LysoPC 6h after tMCAo in plasma, suggesting a higher risk of stroke recurrence.

Cardiolipin is a major GP in mitochondria, especially in the inner membrane, which affects the stability and activity of various membrane protein complexes and metabolite carriers. Our results agree with previous studies showing that after brain injury, plasma cardiolipin is a marker of early brain damage, which corresponds to early neurologic injury and subsequent outcome ^26^. We hypothesize that cardiolipins, once I/R injury occurs, may be released into systemic circulation early after brain mitochondrial dysfunction, where they act as danger signals to exacerbate ischemia damage.

Sphingomyelin is a key component of lipid rafts that play a key role in myelination and the maintenance of myelin ^27^. Given the relative abundance of myelin lipids to axonal protein components, leakage of these lipids into the periphery after brain injury may be a sensitive indicator of acute injury. Stress stimuli, such as acute systemic inflammation, or the release of pro-inflammatory cytokines such as TNF-α or IL-1β, activate the sphingomyelinase pathway. Sphingomyelinase degrades sphingomyelin to ceramide, explaining why 6h after tMCAo we saw a reduction of sphingomyelins and an increase in ceramides or hexosyl ceramides in plasma.

We demonstrated that RIPerC and RIPostC had a brain metabolome signature expressing an adaptive metabolic response with a potential neuroprotective role. In the brain, few metabolites showed significant differences between the three groups. Concretely, two SPs and one GL. RIPerC and RIPostC effects were inverse, as RIPerC decreased ceramide levels while RIPostC increased them. It is known that ceramides may be associated with cell apoptosis ^28^, which may explain the RIPerC effect. Nevertheless, it is common knowledge that very long chain ceramides promote cell proliferation and play a protective role against apoptosis ^29^, which could be the case with the RIPostC effect. Finally, RIPerC increase TG(28:0) levels in the brain, which is a GL that is responsible for storing energy. This long-chain fatty acid is most likely used by the brain as a fuel for the mitochondrial energy generation.

At 6h after tMCAo, both treatments increased oxidized glutathione (GSH) plasma levels. GSH is a scavenger of ROS, indicating that RIC acts as an antioxidant mechanism ^30^. Also, it is known that the brain content of GSH is depleted during I/R. Our results indicate that GSH might inhibit the effects of cerebral infarction and boost antiapoptotic signaling after ischemic stroke, suggesting that GSH may be a potent therapeutic antioxidant that can attenuate stroke injury.

PG(32:2) is a phosphatidylglycerol that was elevated in the plasma 6h after tMCAo in both RIC treatments. It has been described that this compound suppresses inflammation ^31^. Furthermore, ganglioside GM1 was increased. It is well known that this sphingolipid exerts neuroprotective functions ^32^. Also, both RIC strategies increase the levels of LysoPe(18:0), PS(44:4) and CDP-DG(50:0) at 6 hours after tMCAo.

At the same time, RIPerC reduced plasma LysoSM(18:0) and L-linoleoyl glutamine levels, indicating less inflammation, less ROS production, and decreased release of proinflammatory cytokines. On the other hand, RIPostC reduced N-eicosapentaenoyl glutamine, which is related to excitotoxicity, meaning that there was a reduction of the neuronal death process. Moreover, it increased choline levels. Choline is an organonitrogen compound that has neuroprotective roles after stroke ^33^. Choline supplementation increases neuroplasticity and recovery after stroke ^34^. RIPostC also increased DG(34:0) a lipid second messenger.

Seventy-two hours after stroke, RIPerC and RIPostC reduced L-tryptophan levels in plasma. Stroke events are characterized by the augmentation of tryptophan metabolism in the CNS and periphery ^35^. Also, it is a player in inflammation and the immune response. This reduction may explain the reduction in the inflammatory response when RIC was applied. Also, both treatments reduced the benzoid 2,4-Dihydroxybenzophenon (2,4OH-BP). It is known that 2,4OH-BP passes through the BBB, increases the levels of extracellular glutamate, and induces apoptotic processes ^36^. So, the observed decrease may be related to the reduction of apoptotic cells.

It is important to note that both RIC treatments reduced plasma levels of oxidized acylcarnitines, oxidized LysoPC, oxidized TAGs and oxidized PC, suggesting that there was less oxidation of fatty acids and oxidative stress. Also, they reduced sulfatide levels, indicating a reduction of neuronal apoptosis. Moreover, RIPerC and RIPostC increased PA(32:1) levels, which is a precursor of CDP-DAG and DAG that have an important role in cellular signaling and membrane dynamics.

RIPostC reduced hypoxanthine, too. Hypoxanthine is a breakdown product of AMP and increases its concentration in situations of oxygen limitation ^37^. The accumulation of hypoxanthine is a potential source of oxygen free radicals and indicates a disruption in purine metabolism.

Ganglioside GM3 was increased when RIPerC was applied. An increase in ganglioside GM3 expression has been described in both human and animal models of Alzheimer’s disease, as well as in rodent models of stroke, suggesting the activation of an adaptive response from brain injury to offer precursors for the biosynthesis of more complex gangliosides such as GM1, which have neuroprotective effects to preserve neuronal function ^38^.

## CONCLUSION

In conclusion, RIC activates an adaptive response based on increasing circulation levels of structural and bioactive lipids to facilitate functional recovery after brain ischemia. Furthermore, changes after RIC application seems to facilitate the production of anti-inflammatory mediators and to reduce the inflammatory response, contributing to a better immune response. Our study provides first-time evidence of a metabolomic-lipidomic signature related to the development of brain tolerance in tMCAo mice induced by RIPerC and RIPostC and defined by specific regulation of lipid metabolism. Our findings open the way for the future discovery of inflammatory biomarkers and metabolomics targets for new therapies of neuroprotection.

## Acknowledgements.

CTQ received predoctoral fellowships in the Health Research Training program (PFIS contracts) of the Instituto de Salud Carlos III (ISCIII), AES2018 (FI18/00319).

## Authors contributions

Conceived the study (CTQ, GA, RP, FP); designed experiments (CTQ, GA, RP, FP); participated on data interpretation and draft the manuscript (CTQ, GA, RP, FP). All authors critically revised the final version of the manuscript. All authors approved the final version to be published. FP procured funding.

## Data Availability Statement

Requests for access to the data reported in this paper will be considered by the Lead contact on reasonable basis.

---

**Supplementary Figure 1:**
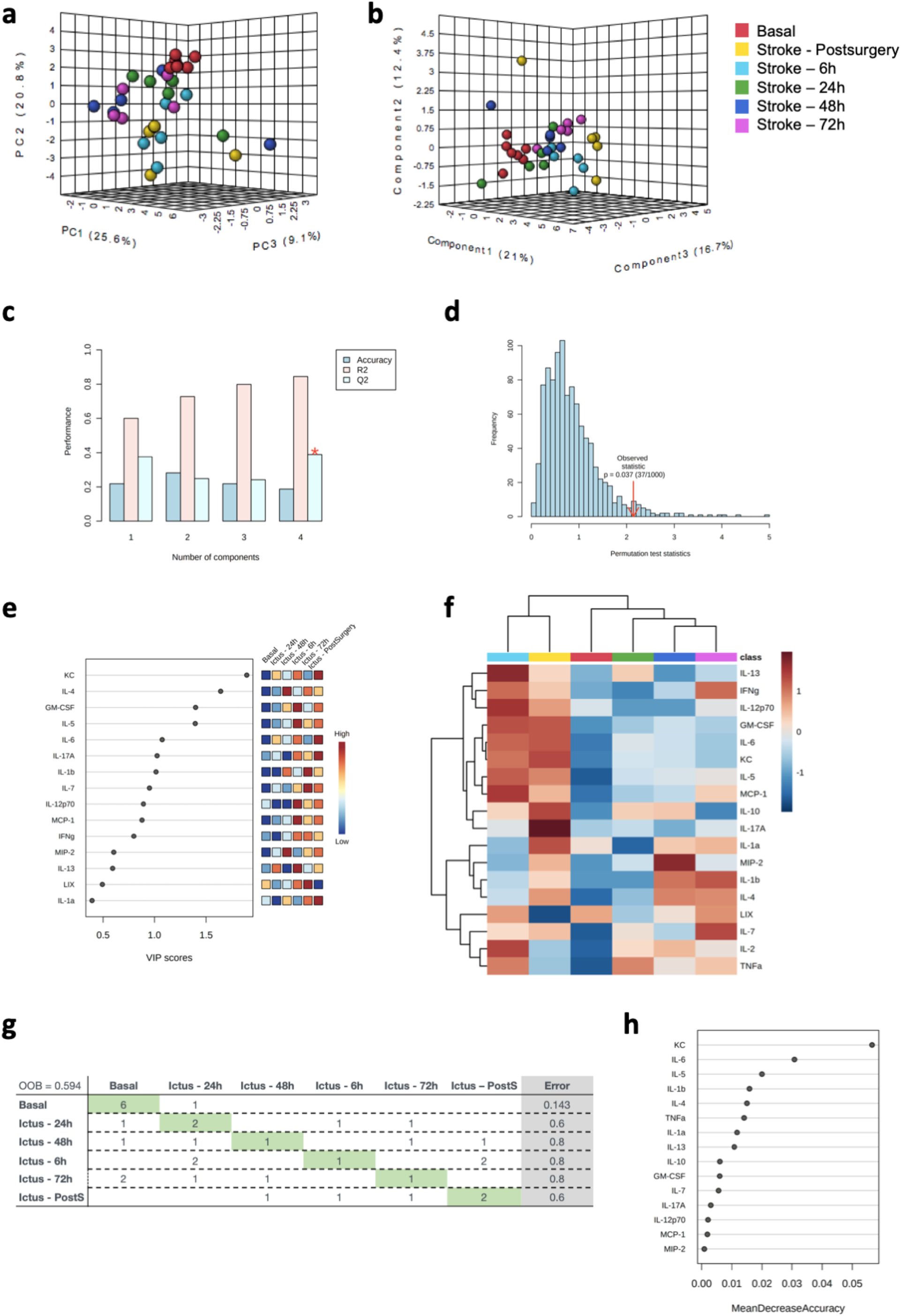
Multivariate statistics reveals a time-specific plasma cytokine profile. **a** Principal component analysis (PCA) representation of cytokine levels of tMCAo mice model. X: Principal component 1 (PC1); Y: Principal component 2 (PC2); Z: Principal component 3 (PC3). **b** Partial least squares discriminant analysis (PLS-DA) representation of cytokine levels of tMCAo mice model. **c** Cross validation (CV) analyses (LOOCV method) of the PLS-DA model). **d** Permutation test (1000 repeats) using separation distance. **e** Variable importance projection (VIP) scores, indicating the elements which contribute the most to define the first component of a PLS-DA. **f** Hierarchical clustering of time points according to average sample values of cytokine abundance of tMCAo mice model. **g** Random Forest (RF) classification algorithm. **h** VIP scores for RF.

**Supplementary Figure 2:**
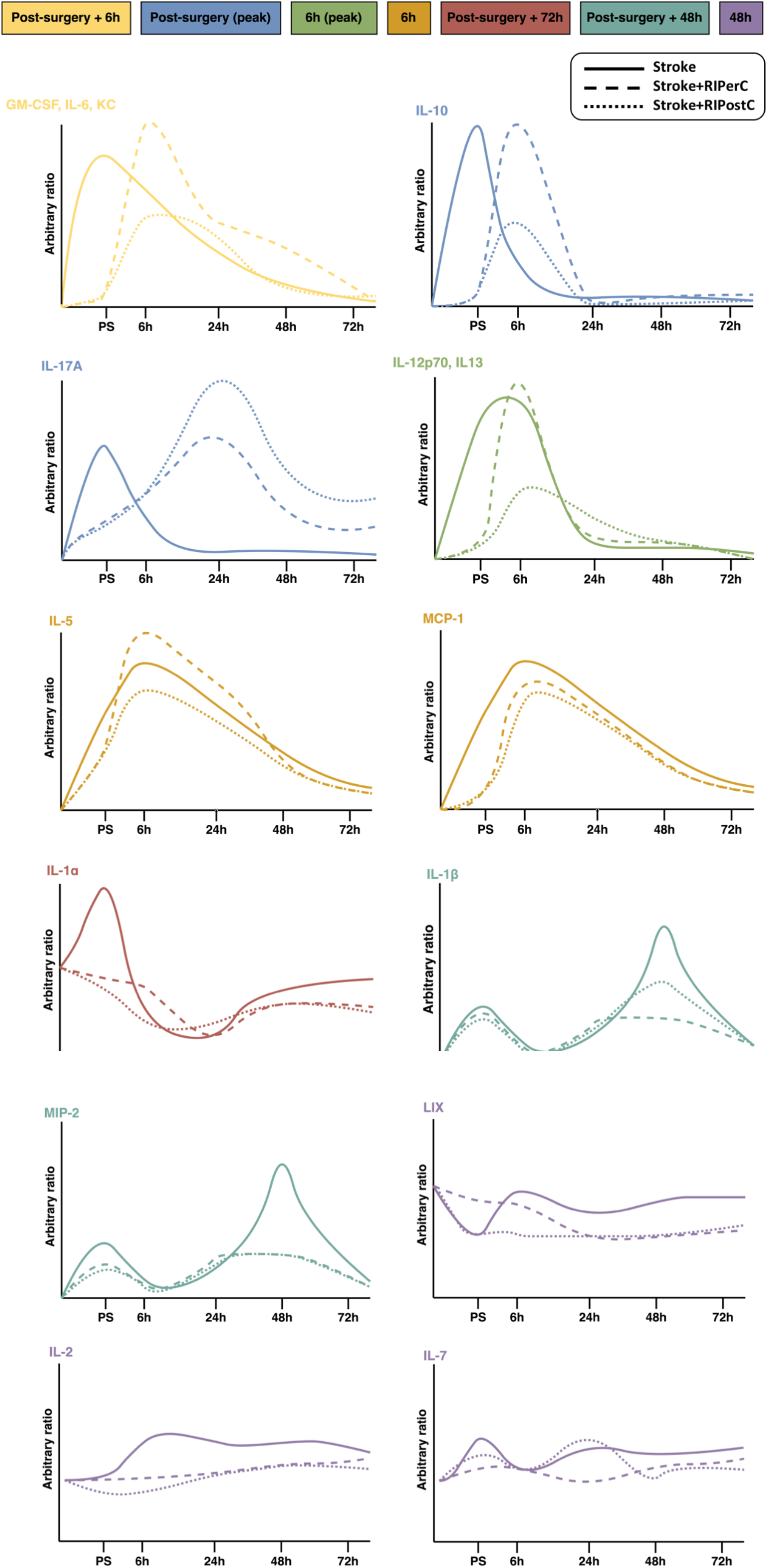
Schematic diagrams summarizing the post-ischemic expression profiles of all analyzed cytokines and inflammatory mediators.

**Supplementary Figure 3:**
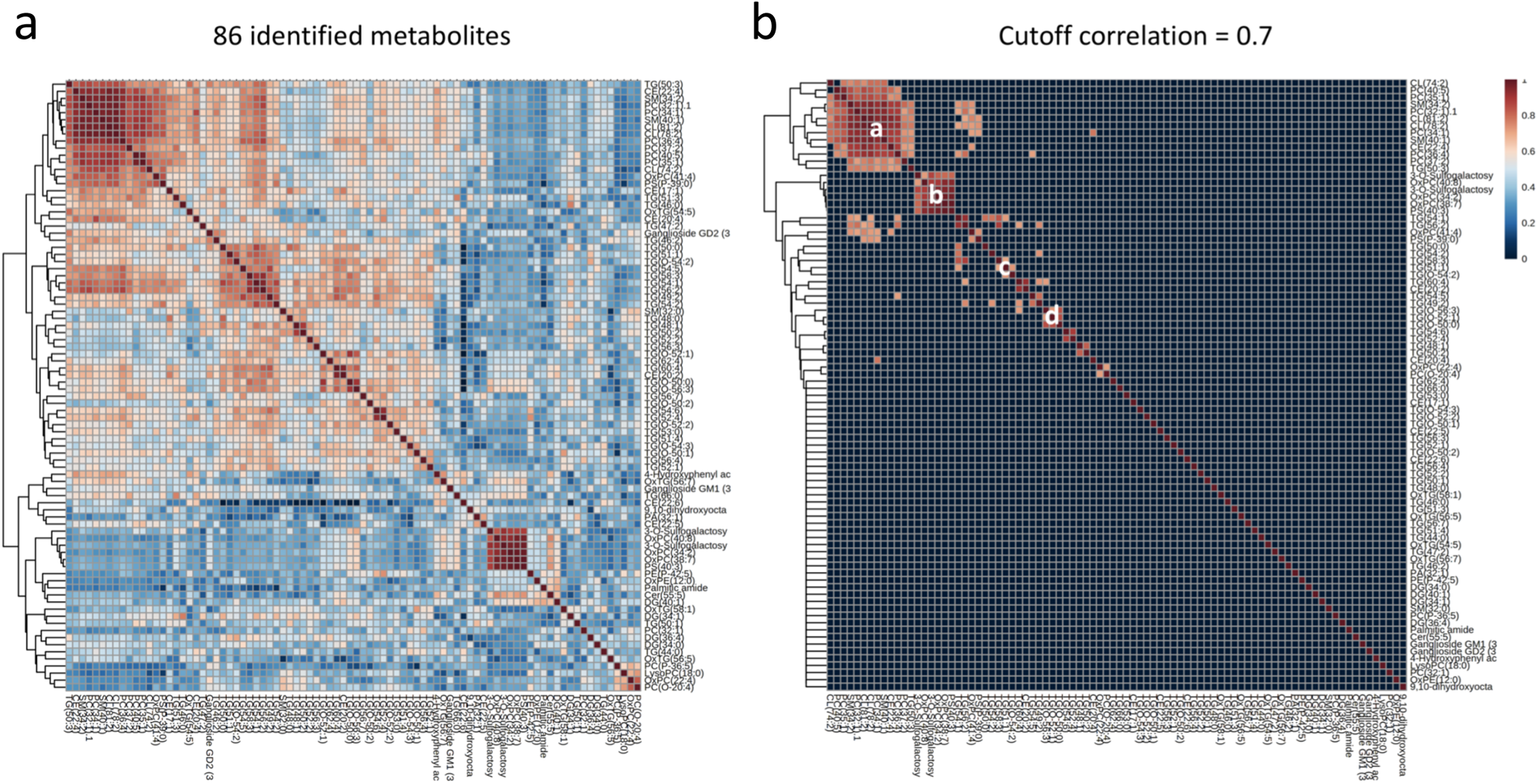
Pearson correlation matrix of identified metabolites represented as a heatmap. **a** Heatmap of Pearson correlation matrix across the 86 identified metabolites with corresponding hierarchical tree. b Heatmap of Pearson correlation matrix across the 86 identified metabolites after selecting a cutoff correlation of 0.7 and p<0.05.

**Supplementary Table 1:** Main source of secretion and function of each cytokine. Adapted from Chen et al., 2018; Nguyen et al., 2016.

